# Thriving to surviving: light wavelength modulates photoacclimation response in the siphonous green alga *Derbesia*

**DOI:** 10.64898/2026.06.07.730690

**Authors:** Riyad Hossen, Saelin Bjornson, Joseph Pelle, John A West, Trevor Bringloe, Kshitij Tandon, Pranali Deore, Heroen Verbruggen

## Abstract

Algae require specific acclimation strategies to cope with spectral variability in shallow marine habitats. We investigated how the siphonous green alga *Derbesia* alters its photosynthetic and metabolic processes under white (WL), blue (BL), green (GL), red (RL), and far-red light (FL) by conducting photobiological and transcriptomic sampling over a 10-day period. Our results show two contrasting photoacclimation strategies: BL and GL promoted metabolic activity associated with growth, whereas FL and RL induced a low-light-like survival strategy characterized by reduced growth and suppression of the core metabolism. Photosynthetic acclimation across all conditions primarily occurs within the light dependent reactions. BL and GL promoted early acclimation marked by the immediate activation of light-harvesting complexes (LHCs) and a key transcriptional regulator *MYB*, and showed better acclimation marked by the sustained activation of ATPases, ATP transporters, and hormone-signaling components. BL induced a distinct transcriptional shift during the transition to prolonged exposure, including enhanced cyclic electron transport, and key regulators of protein synthesis, DNA replication, and transcriptional regulation. In contrast, FL, and to a lesser extent RL, triggered responses resembling low light acclimation with constrained growth, characterized by inefficient energy utilization, enlarged antenna systems, chloroplast proliferation with aggregations, and reduced growth rates. This study suggests high accumulation of core photopigments and reduction in chlorophyll *a*/*b* is an acclimatory response to FL, and consistently higher activation of core metabolic processes under WL likely indicates the evolutionary adaptation of *Derbesia* to shallow coastal environments where broad-spectrum light predominates. Additionally, our newly sequenced draft genome of the *Derbesia* strain for this study could serve as a genomic resource for future molecular photobiology research in Bryopsidales algae.

## INTRODUCTION

Light drives sustenance of life on earth through photosynthesis, and act as a critical signal that regulates growth and development in a range of phototrophs (Chen *et al*., 2004; Izawa *et al*., 2006; Yang & Weathers, 2015; Lee *et al*., 2019). The solar spectrum ranges from ultraviolet to near-infrared (Grant, 1997; Battle *et al*., 2020) of which only photosynthetically active radiation (PAR) ranging from 400 to 700 nm is utilised by plants, majority of algae, and cyanobacterial species for oxygenic photosynthesis (Zhu *et al*., 2010; Chen & Blankenship, 2011). However, in aquatic environments PAR availability is often limited and significantly different compared to land (Gordon, 1989; Kirk, 2011).

In aquatic ecosystems, light availability decreases exponentially with depth of the water column (Gordon, 1989). The light attenuation is not uniform across all wavelengths of PAR; rather, water selectively absorbs certain wavelengths more than others. This effectively transforms the water column into a monochromator-like system that narrows the spectrum of available light at increasing depths (Jerlov, 1976; Falkowski *et al*., 1990). Water rapidly absorbs long-wavelengths such as the red spectrum of light (600–700 nm) within the first few meters, whereas blue (400–500 nm) penetrate deepest in the clear oceanic water due to the absorption characteristics of the water itself. In more turbid coastal waters that are composed of optically active constituents like phytoplankton and colored dissolved organic matters, light shifts to green (500–600 nm) at depth, as these constituents absorb the blue wavelengths (Morel & Prieur, 1977; Kirk, 2011). The far-red spectrum (700–790 nm), once considered photosynthetically irrelevant, is now recognized for its role in supporting photosynthesis in shallow-water habitats shaded by other organisms, such as within dense algal communities, or in the skeletons of reef corals, where upper-layer organisms absorb most of the visible light leaving primarily far-red wavelengths to penetrate into deeper layers (Koehne *et al*., 1999; Kühl *et al*., 2005; Wilhelm & Jakob, 2006; Magnusson *et al*., 2007). These underwater light spectral shifts are critically important for aquatic photosynthetic organisms such as algae, shaping their distribution, physiology, and acclimation strategies (López-Figueroa *et al*., 2003; Terashima *et al*., 2009; Mass *et al*., 2010; Hogewoning *et al*., 2012; Kula *et al*., 2014; Smith *et al*., 2017; Zhen *et al*., 2019).

The Bryopsidales, an order of siphonous green seaweeds, plays a vital role as a primary producer in marine coastal ecosystems and occupies a wide range of ecological niches from intertidal rocky shores to subtidal reef systems (Neil, 1985; Chapman, 1998; Clifton & Clifton, 1999; Kerswell, 2006; Hurd *et al*., 2014). These environments are often characterized by dynamic changes in water depth primarily driven by tidal movements and seasonal variations, thus causing fluctuations in light conditions, particularly the dynamic shifts in light wavelength. These changes pose significant challenges to algal photosynthetic efficiency (Geider *et al*., 2001; Simionato *et al*., 2011; Hurd *et al*., 2014). Many algae employ chromatic photoacclimation, a strategy involving sensing light spectra and regulating pigment composition and light-harvesting machinery to optimize light absorption under shifting spectral conditions (Mass *et al*., 2007; Duanmu *et al*., 2014; Schulze *et al*., 2014). Most of our understanding of these mechanisms comes from studies on a few microalgal lineages (Croce & van Amerongen, 2014; Meneghesso *et al*., 2016; Polukhina *et al*., 2016; Tomas *et al*., 2019). However, research remains limited for green algae, particularly those in the order Bryopsidales. This gap in knowledge is especially notable given the increasing ecological and economic relevance of this group (Verbruggen *et al*., 2009; Lapointe *et al*., 2010; Santos *et al*., 2015; Meinita *et al*., 2022).

Bryopsidales possess a distinctive pigment composition characterized by the keto-carotenoids siphonaxanthin and siphonein (Anderson, 1983; Nakayama & Okada, 1990; Wang *et al*., 2013), that function as major light-harvesting pigments for the absorption of blue-green wavelengths (Raniello *et al*., 2006; Akimoto *et al*., 2007; Marques *et al*., 2021; Seki *et al*., 2022). Experimental studies investigating how light spectrum affects pigment composition, or their functional roles particularly in relation to photosynthetic performance, algal growth, and development in Bryopsidales are scarce. Among few well studied members of Bryopsidales are *Codium tomentosum*, *Caulerpa lentillifera*, *Ostreobium* which have shown diverse but distinct responses in the presence of red light compared to green algae (Lubián *et al*., 2000; Mouget *et al*., 2004; Kang *et al*., 2020; Marques *et al*., 2021; Marques *et al*., 2021). The latter is reported to utilise far-red light for its photosynthesis which is hypothesised to be facilitated by structural modifications of chlorophyll (Chl) *a* along with a reduction of the Chl *a*/*b* ratio (Koehne *et al*., 1999; Wilhelm & Jakob, 2006; Kotabová *et al*., 2014). However, it remains unclear whether this specific acclimation in response to red light occurs in other members of Bryopsidales.

Much of the current understanding is limited to chromatic photoacclimation—the tuning of pigment composition to ambient light quality—while broader aspects of photophysiological adjustment in Bryopsidales, and green algae more broadly, remain underexplored. Beyond its role in photosynthesis, light also acts as a regulatory signal influencing morphogenesis and multiple physiological processes (Franklin, 1998; Mehmood *et al*., 2026). In recent years, there has been growing interest in uncovering the molecular basis of photo-acclimation strategies in algae, particularly through the functional characterization of associated genes (Patelou *et al*., 2020; Thien *et al*., 2021; Obando-Montoya *et al*., 2022). When exposed to different light spectra, algae are known to initiate different signaling cascades and modulate gene expression to adjust their cellular processes, ultimately influencing algal growth and development (Strzepek & Harrison, 2004; Depauw *et al*., 2012; Kim *et al*., 2014). Advances in transcriptomic approaches have played a crucial role in uncovering the genetic underpinnings of these photophysiological responses. Such transcriptome-based insights into light-driven adaptations, are to our knowledge not available for any siphonous green algae, leaving a significant gap in our understanding of their acclimation strategies at the molecular level.

The goal of our study was to investigate the photophysiological responses of the siphonous green alga *Derbesia* sp. to diverse spectral niches, particularly under five distinct spectral conditions (white, blue, green, red, and far-red). By integrating data on growth, chloroplast morphology, light-harvesting pigment composition, Chl *a* fluorescence-based photosynthesis, and transcriptomic profiling, we aim to provide a comprehensive overview of how this alga responds to spectral shifts. We also aimed to obtain a draft nuclear genome sequence of this species to facilitate our analyses and enhance future work.

## METHODS AND MATERIALS

### Algal strain *Derbesia* sp. West4838

The *Derbesia* sp. West4838, belonging to an undescribed species, was originally isolated from Noosa Heads in southern Queensland, Australia in 2013 and has been maintained in culture conditions for 12 years. Species of *Derbesia* are small and morphologically simple representatives of Bryopsidales, convenient for laboratory experimentation. The cultures were monoalgal and maintained in near-axenic conditions following initial antibacterial, antifungal and antidiatom treatments (carbenicillin 0.72 mg mL^-1^; kanamycin 0.03 mg mL^-1^; penicillin G 0.72 mg mL^-1^; chloramphenicol 0.03 mg mL^-1^; GeO_2_ 0.1 mg mL^-1^). Cultures were routinely maintained in filtered seawater enriched with Provasoli nutrient medium and incubated at 26°C under 12:12 h light–dark cycle with an irradiance of 30 µmol photons m⁻² s⁻¹. Approximately half of the media volume was refreshed every 10 days.

### Experimental design

Equal amounts of algal biomass were inoculated in 5 mL of medium in 6-well culturing plates (Greiner bio-one, Kremsmünster, Austria). Cultures were exposed to five different light spectral conditions: far-red (FL, peak at ∼770 nm), red (RL, peak at ∼640 nm), green (GL, peak at ∼550), blue (BL, peak at ∼440 nm), and white LED light (WL, dominant peak at ∼440, another broad peak at ∼530 nm) as shown in Figure 1 for 10 days. Unless mentioned otherwise, all the cultures were maintained at 26°C under 12:12 h light–dark cycle with an irradiance of 30 µmol photons m⁻² s⁻¹. Sampling was performed on the fourth and tenth day from the day of inoculation for growth rate (biological replicates, n=5), photopigment measurements, RNA sequencing, chloroplast morphology and Chl *a* based fluorescence measurement (biological replicates, n=3).

**Figure 1.**
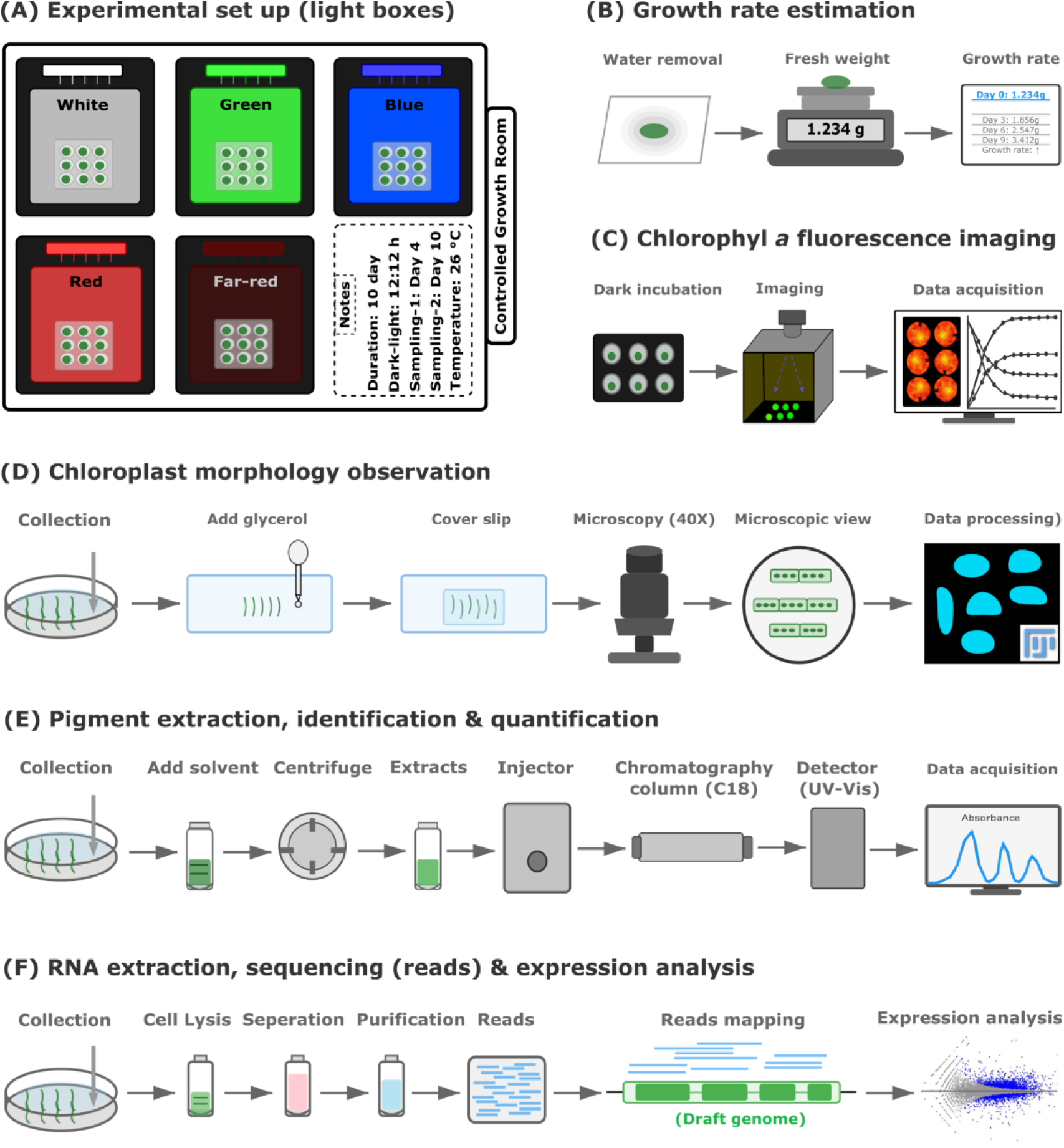
Experimental overview indicating (A) experimental set up, (B) growth rate estimations, (C) chlorophyll *a* based fluorescence measurement, (D) chloroplast morphology observations, (E) photopigments measurement and (F) RNA-seq analysis.

### Chlorophyll *a* fluorescence measurement

Parameters related to photosystem II (PSII) were measured using chlorophyll *a* based fluorescence with imaging-PAM (iPAM, maxi version with RGB head; Heinz Walz GmbH, Germany). The detailed protocols for induction and recovery kinetics, and rapid light curves (RLCs) are provided in supplementary file (method S1). Briefly, algal samples were dark acclimated for 20 minutes prior to measurements. Photosynthetic parameters were obtained using standard saturating pulse protocols on the fourth and tenth day of the experiment. Key parameters measured included: maximum quantum yield of PSII (F_v_/F_m_), effective quantum yield ΦPSII, non-photochemical quenching (NPQ), fraction of open reaction centers (qP), quantum yield of regulated energy dissipation Y(NPQ), non-regulated energy dissipation Y(NO), slow relaxing component of NPQ (qI), state transitions (qT_1_), coefficient of non-photochemical quenching (qN), fast relaxing component of NPQ open also called as energy-dependent quenching (qE), light harvesting efficiency (α), light saturation point (E_k_), and maximum rate of photosynthesis (P_max_). The formulae for these parameters are given in supplementary table S1. All fluorescence measurements started at 10:00 AM, i.e., four hours into the light phase.

### Growth rate and chloroplast morphology estimations

Growth rates were determined by measuring fresh wet biomass. After 10 days, samples were harvested by carefully removing the culture medium through a short-cycle centrifugation step (2000 rpm for 1 min), followed by paper blotting to eliminate excess surface moisture. The final fresh weight of each sample was measured using a precision scale (Mettler Toledo, SN: C017344871). Growth rates were calculated using the formula:

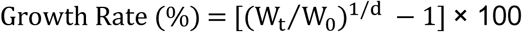

where W₀ and Wₜ represent the initial and final biomass, respectively, and d is the duration of the experiment in days (Yong *et al*., 2013; Ross *et al*., 2017). Chloroplast morphology was examined using a light microscope (Leica, DM750) with 40X objective, equipped with a Canon EOS 600D camera. The acquired images were further analyzed using Fiji distribution of ImageJ (version 1.54f; Schindelin *et al*., 2012) to quantify chloroplast density (via total number of chloroplasts/total area of filament), mean surface area, and shape of each chloroplast. All images were converted to 8-bit grayscale and subjected to background subtraction. For segmentation of individual chloroplast in an image, auto-thresholding was applied followed by a watershed binary process. The segmentation mask was used to count numbers of chloroplast, and estimate mean surface area, size and shape.

### Photopigment analysis

Approximately 4 mg of freeze-dried (−80°C) *Derbesia* biomass was added in centrifuge tubes containing 1.5 mL of ice-cold extraction solvent (95% methanol mixed with 2% ammonium acetate, v/v). All samples were sonicated for 30 seconds at 4°C, followed by incubation at −20°C for 20 minutes in the dark. The extracts were centrifuged at 1100 x *g* for 5 minutes at 4°C, and supernatants were filtered through 0.45 µm syringe filters. A total of 500 µL of the filtered extract was injected into the ultra high performance liquid chromatography (UHPLC, Nexera X2 System, with a diode array detector, SPD-M30A, Shimadzu) located at the HMSTrust Lab, Monash University.

The chromatographic separation of photopigment extracts were performed using a method described by (Sommella *et al*., 2018), with minor modifications. Briefly, 6 µL of samples were injected to a reverse phase C18 column (InfinityLab Poroshell 120 EC-C18, 2.1 × 50 mm, 2.7 µm, Agilent) using a gradient of solvent A (10 mM ammonium acetate, w/v, in Milli-Q water) and solvent B (60:25:15, v/v, acetonitrile:isopropanol:methanol) while maintaining the column oven temperature at 40°C. Gradient for the chromatographic run was started at 60% of B for 1 minute, followed by an increase to 100% of B over 4 minutes, then 2 minutes at 100% of B, followed by a 0.30 minute gradient back to 60% of B, and the 1 minute at 60% of B, with a flow rate of 0.4 mL min⁻¹. The sampling speed was set to 5 µL s⁻¹ with an acquisition scan rate of 12.5 Hz. All photopigments were profiled across the UV-visible range (200–700 nm). The peaks were identified using extracted ion chromatograms at 430 nm, considering primary (absorption maxima) and secondary peaks for each UV profile. All the peaks were further compared to retention times and absorption spectra of commercially available reference standards (Chl *a*, *b*, zeaxanthin, lutein, neoxanthin, fucoxanthin from Sigma-Aldrich) prepared in 95% methanol (v/v in milliQ), and with previously published works (supplementary table S2).

### RNA-seq analysis

Samples were flash-frozen in liquid nitrogen immediately after collection on the fourth and tenth days of the experiment, and total RNA was extracted using the Qiagen RNeasy Plant Mini Kit, following the manufacturer’s protocol. The extracted RNA was then dried using a SpeedVac and stored at −80°C until sequencing. After re-hydration, sequencing libraries were prepared with the VAHTS Universal RNA library preparation kit with poly-A selection, and sequencing was performed on the Illumina NovaSeq platform with 150 bp paired-end reads.

Details of raw RNA reads post-processing, transcriptome assembly, and downstream post-processing analyses followed the methods described in Hossen *et al*. (2026). Briefly raw RNA reads underwent quality assessment with FASTQC v0.12.1, followed by adapter trimming and low-quality read filtering using Trimmomatic (parameters: removal of reads with Phred scores < 20, adapter trimming with Cutadapt) (Bolger *et al*., 2014). A second round of FASTQC quality checks was performed, and results were summarized using MultiQC. To retain only mRNA sequences, SortMeRNA was used with the SILVA rRNA database for ribosomal RNA removal (Kopylova *et al*., 2012). Filtered reads were then aligned to the reference genome (described in the following section) using HISAT2 with default parameters optimized for paired-end reads.

Gene expression was quantified using the FeatureCounts tool (parameters: paired-end mode, counting fragments as single units, excluding chimeric fragments, with default settings for all other parameters) (Liao *et al*., 2014). A count matrix was generated using the “Generate Count Matrix” tool, and differential gene expression (DEG) analysis was performed using the DESeq2 package in R (Love *et al*., 2014). DESeq2 normalization was applied using the default geometric mean-based approach, and differentially expressed genes (DEGs) were identified using a log₂ fold-change threshold of ≥1 and an adjusted p-value ≤0.05.

The DEGs were analysed using two complementary approaches to capture both the temporal dynamics of gene expression within an individual treatment, and treatment-specific responses relative to white light. First, temporal comparisons were conducted within each individual treatment to examine how gene expression changed over time during prolonged exposure. In this analysis, gene expressions at day 10 was compared with day 4 within the same light condition (e.g., BL day 10 vs. BL day 4). This approach allowed us to visualise the temporal progression of transcriptional responses over time within each treatment. Second, treatment-specific responses were assessed by comparing each monochromatic light treatment (BL, GL, RL, and FL) against the white light (WL) at each sampling time. DEGs analyses were therefore performed separately for day 4 (e.g., RL vs. WL at day 4) and day 10 (e.g., RL vs. WL at day 10). This approach enabled identification of genes specifically responsive to different light spectral conditions relative to white light.

In addition, to see the relative expression levels of responsive genes across all samples, we performed standardized expression analysis using only the genes that were identified as significant DEGs in the previous analyses. Raw expression values of these genes were log₂-transformed and standardized using Z-score normalization, enabling direct comparison of relative expression patterns among treatments. This analysis was conducted separately for day 4 and day 10 datasets to visualize expression trends of the selected genes across all light conditions.

Functional analyses were performed in R using annotations from the draft reference genome, with KEGG pathway enrichment conducted using the KEGGREST package (Tenenbaum *et al*., 2019). All raw read sequences are submitted to NCBI under the Bioproject PRJNA1250039.

### Draft genome sequencing, assembly and annotations

To support transcriptome analyses and enhance future work on our *Derbesia* strain, we sequenced its draft genome. Genomic DNA was extracted following the protocol described in Hossen *et al*. (2025), and sequencing was performed using the Illumina short-read platform. For long-read sequencing, extraction, we adapted a phenol:chloroform DNA extraction protocol from Nakamura-Gouvea *et al*., 2022. Nanopore libraries were prepared with the Oxford Nanopore Ligation Sequencing Kit (SQK-LSK114) and sequenced on the PromethION platform with FLO-PRO114M flow cells. Raw FAST5 files were base-called with Guppy v6.4.6 software obtained from Oxford Nanopore Technologies, using the High Accuracy (HAC) model.

For short-read sequences, the Illumina platform successfully yielded 60 Gb of data. However, long-read sequencing proved challenging due to pore clogging, and only 62 Mb of Nanopore data with an N50 of 4,409 bp was generated. To best leverage this sparse long-read data, various assembly tools, parameters and workflows were tested, evaluated by comparing missing BUSCOs in the assemblies. The best assembly was found as follows: short reads were assembled with SPAdes (Bankevich *et al*., 2012) using default parameters, and Redundans (Leszek *et al*., 2016) was used on the assembled draft to carry out short read scaffolding, gap-filling and redundancy reduction. Nanopore reads were then used to scaffold the genome with LRScaf (Qin *et al*., 2019), which was followed again by Redundans using short reads. RNA short reads were then used to better scaffold the genome genic regions with P_RNA_scaffolder (Zhu *et al*., 2018). A round of GapCloser (Luo *et al*., 2012) was then employed to fill in remaining gaps with short reads. Polishing of the final assembly was then done using five rounds of Pilon (Walker *et al*., 2014).

To remove contaminating sequences from the assembly, Infernal (Nawrocki & Eddy, 2013) was first used to search for LSU and SSU rRNA sequences. rRNA genes found were then searched against the SILVA database to remove bacterial sequences. Then, contigs over 20kb in length were split into 10kb chunks, and these and contigs < 20kb were searched against UniProt Reference Proteomes database with diamond blastx, and to the NCBI nt database with blastn. Coverage of original contigs was determined by alignment of short reads with Bowtie2 (Langmead *et al*., 2012). Contig database hits, coverage and GC content was used with the BlobToolKit (Challis *et al*., 2020) to remove contigs suspected to be of non-algal origin.

To predict protein-coding genes, the genome was softmasked with WindowMasker (Morgulis *et al*., 2006), Tandem Repeat Finder (Benson, 1999), and RepeatMasker (Chen, 2004), using both Viridiplantae families and families predicted *de novo* in the assembly with RepeatModeler2 (Flynn *et al*., 2020). RNA Illumina reads were aligned to the softmasked genome with Hisat2 (Kim *et al*., 2019). BRAKER1 (Hoff *et al*., 2016) was used to annotate genes from RNA evidence alone. BRAKER2 (Bruna *et al*., 2021) was then used to annotate genes from protein alignments of the Viridiplantae portion of the OrthoDB v11 database. Final transcripts from the two annotations were selected with TSEBRA (Gabriel *et al*., 2021).

Functional annotation was performed using multiple databases, including EggNOG (Viridiplantae), KEGG, InterProScan, UniProtKB, Phytozome (*Chlamydomonas reinhardtii v5.6* and *Arabidopsis thaliana Araport11*). Annotations were primarily assigned based on EggNOG (Viridiplantae) and KEGG, while Phytozome, InterProScan and UniProtKB were used to resolve ambiguous or missing functions. Sequence similarity searches were conducted using NCBI BLAST+ with default parameters (outfmt 6; maximum target sequences = 10). The best hit for each query was selected based on a combination of lowest E-value and highest percent identity, followed by random manual validation against NCBI and UniProt databases to increase annotation confidence.

### Statistical analysis and data visualization

All statistical analyses, including one-way analysis of variance (ANOVA) and Tukey’s post hoc tests, were performed in R version 4.4.0 (R Core Team, 2024) to evaluate differences in photosynthetic parameters, growth, pigment composition, and chloroplast morphology across all spectral conditions. Due to some replicate measurements failing, particularly in Chl *a* fluorescence parameters, Estimated Marginal Means (EMMs) were calculated using the emmeans package (Lenth, 2023); details of the statistical approach applied for photosynthetic parameters are in the supplemental method S2. The level of significance was *p* < 0.05. Data visualization and heatmaps for differentially expressed genes were generated using ggplot2 (Wickham, 2016) and ComplexHeatmap packages, respectively. Inkscape v1.3.2 was used for creating final illustrations.

## RESULTS

### Draft genome of *Derbesia* West4838

The assembled genome of *Derbesia* sp. was 81.2 Mb in length, comprising 13,469 scaffolds with an N50 of 21,138 bp and a GC content of 42%. Short read coverage of the genome was 738.9x. A total of 13,192 genes encoding 20,003 transcript sequences were predicted, and 82% of all genes and 86% of predicted transcripts had RNA read coverage. The draft genome assembly recovered 87.7% of the conserved genes in the Viridiplantae_odb10 BUSCO dataset, comprising 1,289 complete single-copy BUSCOs, 43 complete duplicated BUSCOs, and 7 fragmented BUSCOs. Functional annotation using multiple databases revealed 3,409 KEGG-assigned genes (BlastKOALA), 4,842 GO-annotated genes (InterProScan), and 6,168 genes annotated using the EggNOG Viridiplantae database.

### Transcriptome profiles across light spectral conditions

Transcriptomic responses varied in the number of differentially expressed genes (DEGs) both over time for in-between treatments and compared to white light (WL). Overall expression profiles are broadly resolved in two distinct patterns corresponding to wavelengths, separating the short-wavelength (blue light, BL, and green light, GL) and long-wavelength treatments (far-red light, FL, and red light, RL). Relative expression analysis based on standardized Z-score normalization across all samples reflected this pattern, and showed short-wavelength treatments were generally associated with stronger gene activation at the early stage (Fig.2 & 3). The identified DEGs, including their relative expression by Z-scores are presented below in two broad functional categories.

**Figure 2.**
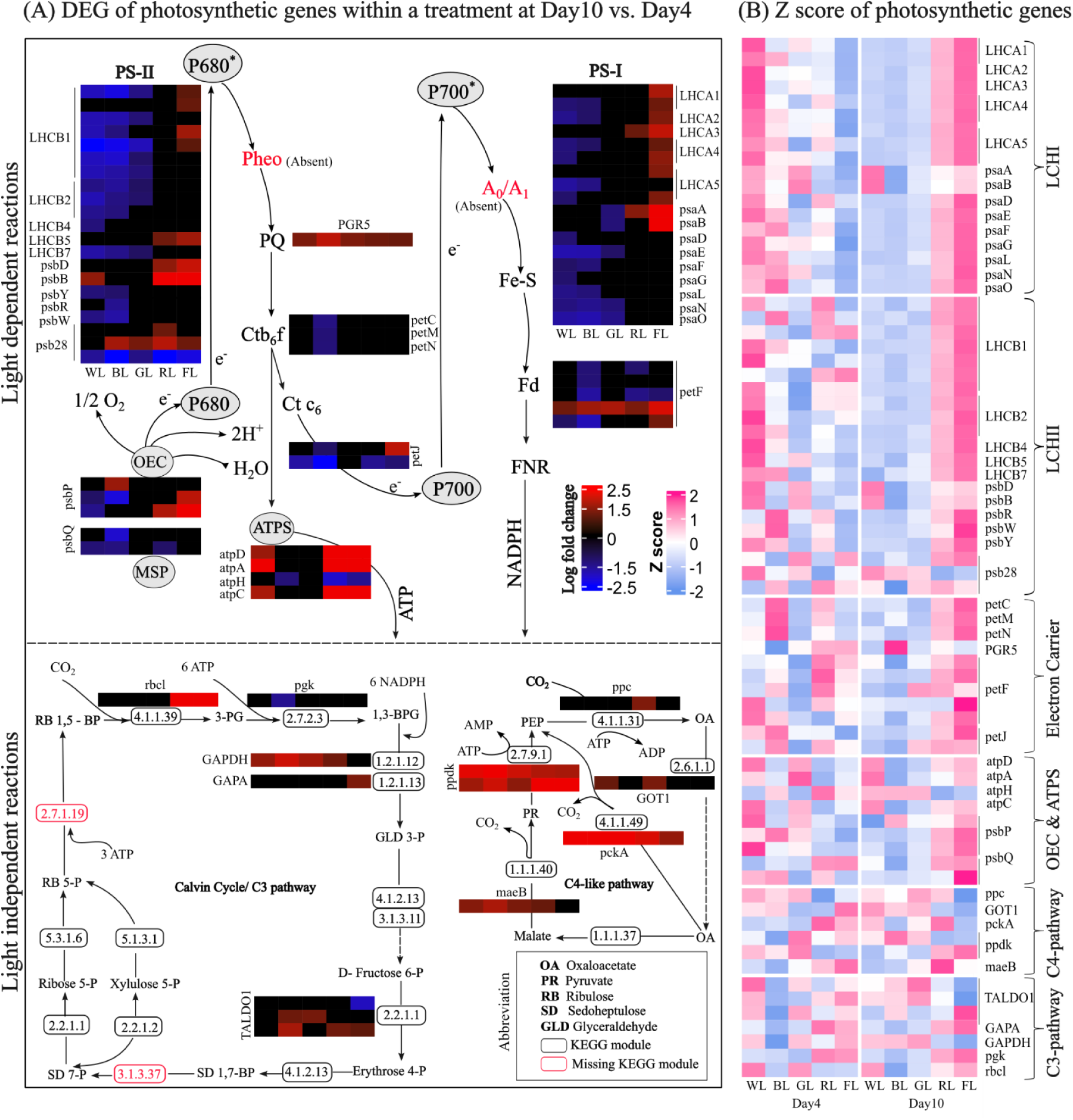
Transcriptomic responses of photosynthesis in response to different light spectra. (A) Differentially expressed genes (DEGs) across time (day 10 vs. day 4) for each light condition (B) Relative expression (Z-score) across all light conditions.

**Figure 3.**
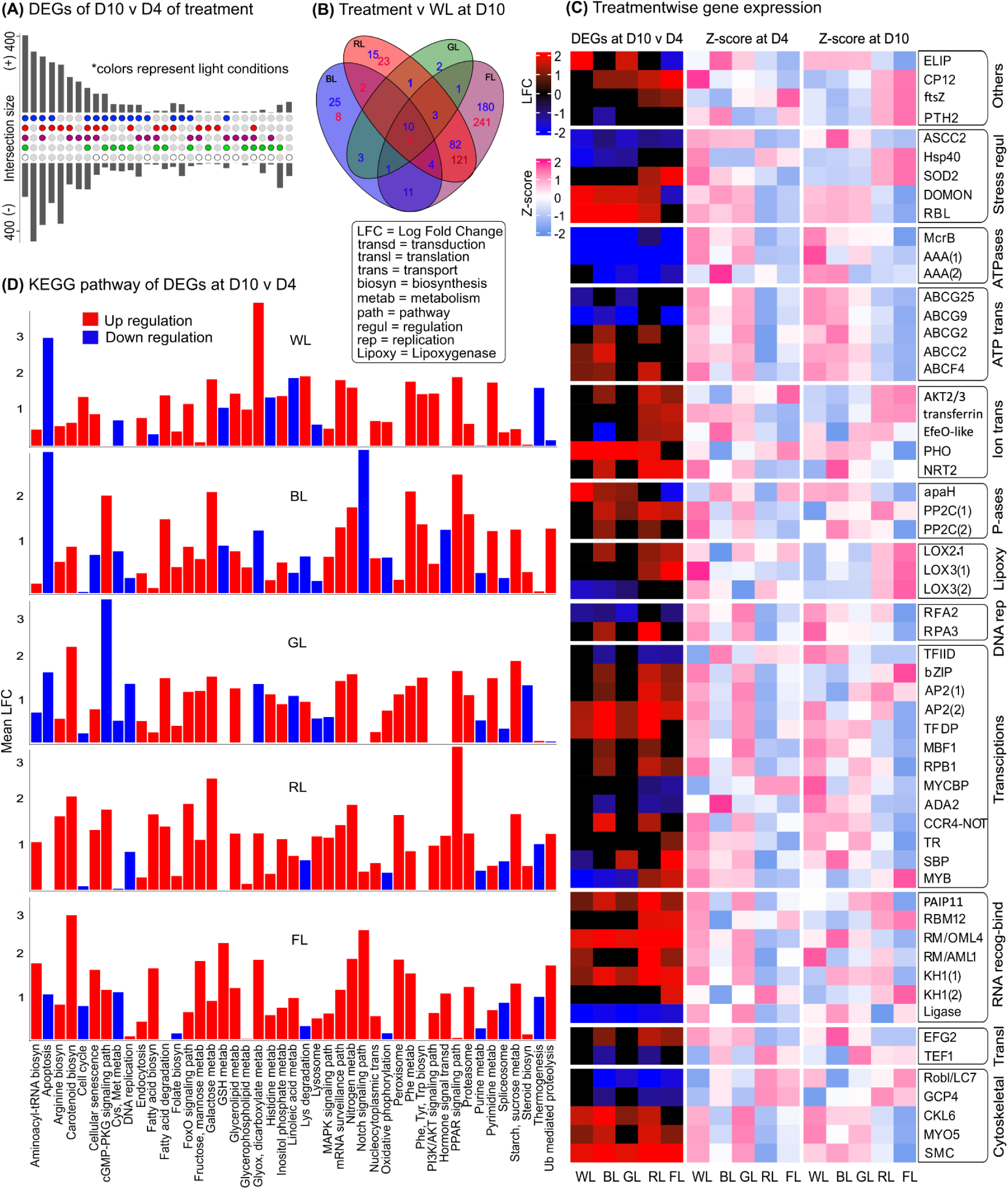
Transcriptomic responses of genes associated with key cellular processes beyond photosynthesis and carbon metabolism. Venn diagrams show the total number and overlap of DEGs across treatments at day 10 vs. day 4 (A), while the UpSet plot compares individual treatments with WL (B). KEGG pathways are shown in (C). Heatmaps depict the expression patterns of representative genes involved in major cellular processes (D).

#### Gene expression of photosynthesis and central carbon metabolism

The DEG and Z-score analyses indicated that treatment-associated transcriptomic responses in photosynthetic processes were related mostly to the light-dependent reactions (Fig. 2). Differences in expression between days 4 and 10 (Fig. 2A) showed downregulation of most genes encoding for light-harvesting complexes (LHCII and some LHCI), except for *psaA* and *psaB* in WL and BL, and to a lesser extent in GL, while upregulated in FL (Fig. 2A). In Z-score analysis, these genes showed relatively higher expression in WL, BL and to lesser extent in GL at day 4, while at day 10, expression of these genes became higher under FL, followed by RL (Fig. 2B). Electron transport chain (ETC) genes, including *petC, petM, petN*, *petJ*, and *petF*, were downregulated over time under BL, consistent with their initially high relative expression. The proton gradient regulation 5 *(PGR5)*-like gene, which is associated with cyclic electron flow, showed markedly higher relative expression at day 10, particularly in BL. Genes associated with ATP synthases (ATPS), except *atpH*, showed upregulation over time in WL, FL and RL, but remained unchanged in BL and GL. These ATPS genes, along with the key photosystem genes (*psaA*, *psaB*, *psbD*, *psbB*), maintained consistently higher relative expression under WL throughout the experiment. FL showed relatively higher expression rates of photosynthesis-related genes on day 10, and the DEGs analysis of FL vs. WL showed that most genes of the light-dependent phase (e.g., *psbP*, *petF*, *atpD*, *psaA*, *psaB*, *LHCA3*, *LHCB1*) were enhanced at day 10 compared to day 4 (Supplementary Fig. S20).

Genes involved in central carbon metabolism remained largely unaffected across treatments, with few exceptions. The marker gene for C3 photosynthetic cycle Ribulose-1,5-bisphosphate carboxylase large subunit (*rbc*L) showed upregulation in FL and RL on day 10, while another marker gene glyceraldehyde-3-phosphate dehydrogenase (*GAPDH*) showed upregulation at WL, BL and GL at day 10. For the C4-like pathway, phosphoenolpyruvate carboxykinase (*pckA*) and malate dehydrogenase (*maeB*) were upregulated in all treatments except FL, where they remained unchanged.

#### Gene expression of other key cellular processes

Z scores showed higher expression of genes for ATPases, ATP transporters, high affinity nitrate transporter 2 (*NRT2*), *apaH*-like phosphatase and RNA-recognition and binding regulators at day 4 under GL, at day 10 under BL, and at both timepoints under WL (Fig. 3C). Likewise, the transcriptional co-activator Multiprotein Bridging Factor 1 (*MBF1*), the global regulator complex for gene expression maintenance (*CCR4–NOT*) and the chloroplast/mitochondrial elongation factor G2 (*EFG2*) showed high expression early in GL, high later in BL, consistently high expression in WL, and consistently low expression in RL and FL. MYC-binding proteins gene (*MYCBP*), on the other hand, showed higher expression early under RL and FL. A key growth regulating transcription factor gene (*MYB*) was most strongly expressed in FL at day 10.

The early light-induced protein (*ELIP*) gene associated with the regulation of chlorophyll biosynthesis, pigments binding, and stabilization of the pigment–protein complexes in thylakoids (Tzvetkova-Chevolleau *et al*., 2007; Liu *et al*., 2020) showed consistently higher expression under BL and GL, but extremely low expression in FL and RL. In contrast, the chloroplast division gene *ftsZ* was strongly expressed throughout FL (and to a lesser extent in RL) and showed very low expression in WL, BL, and GL. Stress-regulation, particularly cell signaling and redox transport genes encoding dopamine β-monooxygenase N-terminal (DOMON)-domain proteins and Rhomboid-like (*RBL*) serine proteases, were consistently highly expressed in WL, BL, and GL, while lowly expressed at FL and RL. Lipoxygenases were highly activated later in FL and to a lesser extent in RL. DEGs based temporal transcriptomic shifts showed that most of these genes’ responses changed over time largely, mostly showing upregulation in the BL, RL and FL conditions.

KEGG pathway analyses showed apoptosis-like programmed cell death highly downregulated in GL and BL over time (Fig. 3D). cGMP–PKG signaling was downregulated in GL, whereas linoleic acid metabolism was downregulated in WL, BL, and GL but upregulated in FL and RL later. When compared to WL, core growth- and metabolism-associated pathways, including hormone signaling, MAPK signaling, nitrogen metabolism, apoptosis, and protein processing in the ER, were strongly downregulated in RL and FL. Moreover, cellular senescence was strongly upregulated in FL, followed by RL, and glutathione metabolism also showed marked upregulation in FL at late stage and similar results found when compared to WL (Supplementary Fig. S8, S9; S11 & S12).

### Chlorophyll *a* fluorescence

We assessed differences in photosynthetic performance by measuring Chl *a* fluorescence, which revealed a clear distinction between light spectral treatments on day 4 (Fig. 4A-D). The temporal changes in the effective quantum yield of PSII (ΦPSII) within each individual light condition followed the same pattern as the observed difference between day 4 and day 10 in our RNA-seq analysis. GL exhibited higher ΦPSII at day 4 than at day 10, whereas BL showed higher ΦPSII later in the experiment. The maximum quantum yield of PSII (F_v_/F_m_) in dark-acclimated samples, maximum photosynthetic rate (P_max_), and light-harvesting efficiency (α) were higher at day 4 in BL and GL than at day 10, while in FL, these parameters increased over time (Fig. 4A,B, D, supplementary Fig. S19). Across WL, GL, and BL, *Derbesia* displayed lower non-photochemical quenching (NPQ), lower slow relaxing NPQ component (qI), and state transition 1 (qT_1_) than observed in FL and RL in both time points. Temporal changes of these parameters showed clear increase at day 10 compared to day 4 in FL and RL. We also observed a small amount of energy-dependent quenching (qE), associated with heat dissipation through pigment cycling, across all light conditions (Fig. 4C).

**Figure 4.**
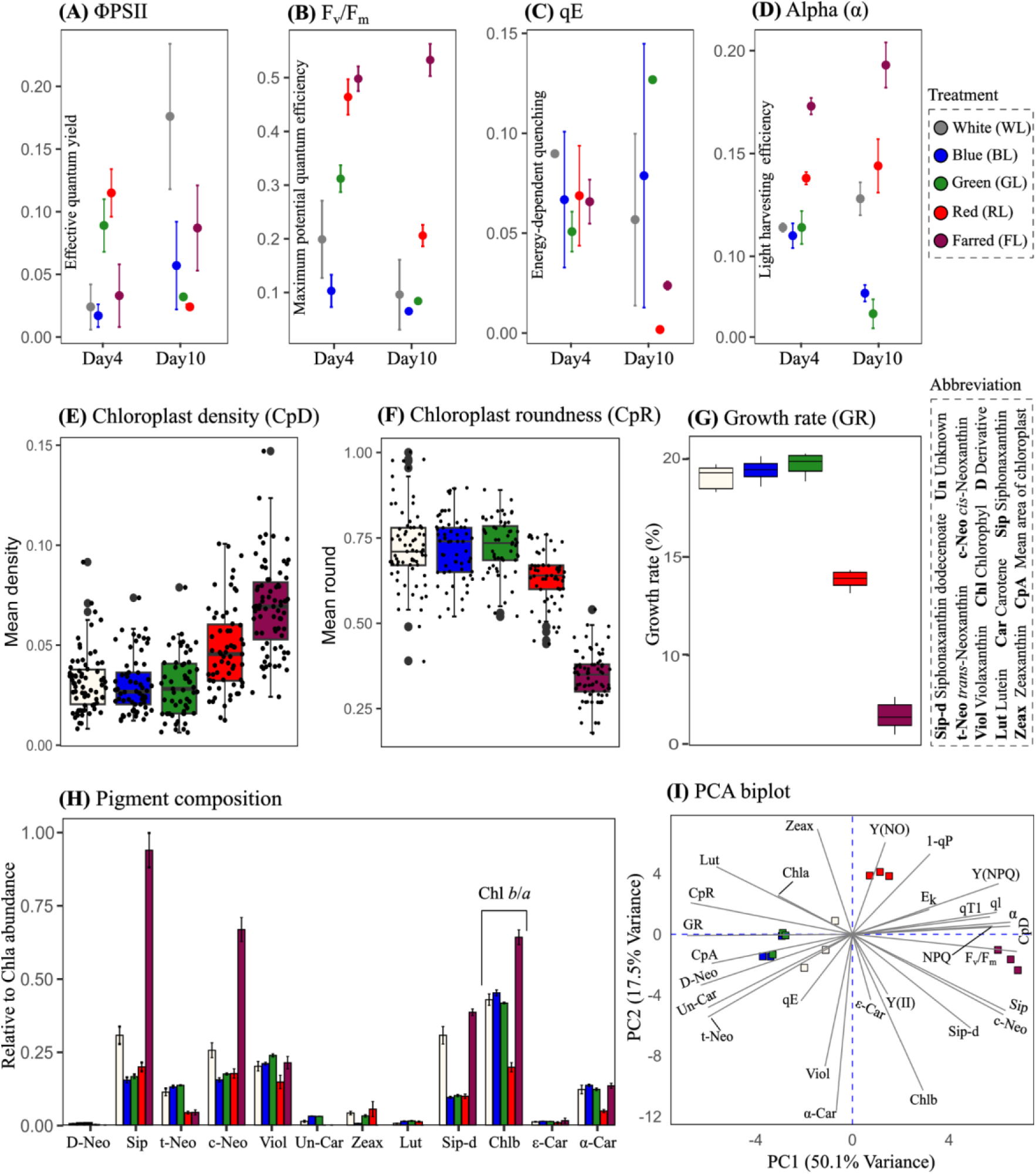
Physiological responses under different light treatments. Shown are chlorophyll *a* fluorescence parameters (A–D), chloroplast morphology (E,F), biomass growth (G), photopigment composition (H), and a PCA bi-plot showing variation across physiological parameters (I). Detailed statistical analyses with level of significance *p* < 0.05 are provided in Supplementary Tables S3–S5.

### Chloroplast morphology, photopigment composition and biomass growth

In WL, GL, and BL, chloroplasts were nearly round with relatively large surface areas, whereas in FL and RL they appeared more elongated and densely arranged within the algal filaments (Fig. 4E, F; Supplementary Fig. S18). FL produced the most pronounced narrowing of chloroplasts, reflected in the lowest mean chloroplast roundness or circularity. Additionally, large and conspicuous pyrenoids were consistently observed in FL-treated chloroplasts (Supplementary S21). Growth rate was highest in GL (19.69%), BL (19.38%), and WL (19.05%), and was markedly reduced in FL (6.94%) (Fig. 4G).

Photopigments analysis revealed that GL, BL, and WL treatments contained the maximum 13 examined pigments, whereas RL lacked the unidentified carotenoid (Un-Car), and FL lacked Un-Car, Zeaxanthin (Zeax), lutein (Lut), and the neoxanthin derivative (D-Neo) (Fig. 4H). The core antenna pigments (i.e., Chl *b*, siphonaxanthin, and *c*-neoxanthin) accumulated most strongly in FL, and the Chl *a*/*b* ratio was lowest in FL. The accumulation of all-*trans*-neoxanthin (*t*-Neo), a pigment proposed to contribute to heat-induced energy dissipation, was observed across the treatments with highest in GL and BL conditions. The principal component analysis (PCA) showed that the *t*-neoxanthin accumulation has a clear positive association with energy-dependent quenching qE (Fig. 4I).

## DISCUSSION

We investigated the effect of different light spectra (blue, green, white, red and far-red) in modulating photosynthetic and metabolic responses of the siphonous green alga, *Derbesia* sp., to better understand its acclimation strategies in different light spectral niches. Our results clearly indicate that *Derbesia* showed two distinct photoacclimation strategies: a rapid acclimation response from the onset of short-wavelength light exposure, and a slower response characteristic of low-light acclimation when exposed to long-wavelength spectral conditions.

### Short wavelength light promotes rapid photosynthetic optimisation and sustained energy production

Our transcriptomic data indicated that in blue (BL) and green (GL) light, photosynthetic adjustment in *Derbesia* occurs primarily through modulation of light-dependent reactions, with light-harvesting complexes (LHCI & LHCII) being particularly responsive. The early upregulation of LHCII genes under both BL and GL suggests blue and green spectra drives photosynthetic activity. The subsequent downregulation of many LHCs genes likely reflects successful early photoacclimation; once the photosynthetic system has adjusted, further production of antenna proteins is reduced, a response often associated with improved light-use efficiency (Melis, 2009; Kirst *et al*., 2018).

This early photoacclimation is also supported by declining trends in Chl *a* fluorescence parameters F_v_/F_m_ and α, indicating functional downscaling of the PSII apparatus later (Henley, 1993; Baker, 2008). BL promoted stronger non-cyclic electron flow during early exposure, whereas the subsequent decline in electron availability was likely compensated by enhanced cyclic electron flow, as indicated by upregulation of *PGR5*-like genes (Suorsa *et al*., 2016; Ma *et al*., 2021). This suggests that in BL, *Derbesia* maintained its electron transport to support sustained ATP synthesis throughout the experiment, in contrast to GL, which showed no clear temporal changes in these downstream light dependent reaction components. However, Chl *a* fluorescence measurement revealed higher ΦPSII early in GL and later in BL, suggesting that green light initially supported more efficient utilization of absorbed light energy by PSII, whereas blue light showed improvement over time.

The early exposure photosynthetic adjustment in BL and GL appears to be supported by a higher supply of ATP-derived energy, reflected by early activation of the genes associated with ATPases. However, the relatively high expression of ATPases under BL and GL compared with other monochromatic light conditions throughout the experiment suggests that *Derbesia* remained metabolically more active under these light conditions. This pattern of acclimation is further supported by consistently higher expression of ATP transporter genes, likely facilitating ATP-driven transmembrane transport of ions, nutrients, lipids, and other metabolites (Kühlbrandt, 2004; Morth *et al*., 2011) to maintain growth and development. Although both BL and GL maintained high activation of ATPases and ATP transporters throughout the experiment, BL comparatively showed stronger expression during prolonged exposure, exceeding its early-stage levels, indicating that *Derbesia* remained more metabolically active under BL during prolonged exposure.

### Sustained metabolic activity under short wavelengths eventually triggers nutrient-limited stress

We also observed consistently higher activation of other key metabolic pathways in BL and GL compared to other monochromatic light conditions. These responses are likely essential for maintaining gene expression, protein synthesis, and stress management with observed higher biomass growth. Temporal DEGs analysis revealed that in BL, and to some extent GL, *Derbesia* employs stage-specific strategies to sustain acclimation over time. Early on, *Derbesia* promotes programmed cell death, DNA replication pathways, and induced *MYB* expression, which all are crucial for maintaining initial growth and development (Mariyam *et al*., 2025). Prolonged acclimation in these light conditions showed induction of *EFG2* and *RPA3,* key regulators for protein synthesis and DNA replication (Kaul *et al*., 2011; Gou *et al*., 2017; Dassi, 2017; He *et al*., 2018), also marked by the induction of *MBF1* which is involved in growth hormone regulation (Tsuda *et al*., 2004).

Despite the activation of growth and development associated pathways throughout the experiment, prolonged exposure also appeared to impose physiological constraints, such as the induction of stress-responsive genes at later stages. This likely resulted from resource limitations, as the experiment was conducted in a closed culture system. Because BL, GL, and WL supported substantially higher growth than RL and FL, these cultures would also have depleted nutrients more rapidly. In particular nitrate availability may have been reduced, reflected in the BL condition by the late-stage induction of high-affinity nitrate transporters, a known indicator of nitrogen limitation (Lezhneva *et al*., 2014; Kiba and Krapp, 2016). In addition, increased activity of *ApaH*-like phosphatases is associated with nutritional limitation and broader metabolic stress responses (Monds *et al*., 2010; Sasaki *et al*., 2014; Zegarra *et al*., 2023). This increased activity of *ApaH*-like phosphatases at later stages further suggests that *Derbesia* was likely regulating its metabolism to cope with nutrient limitation, particularly phosphate limitation, a response previously observed in bacterial cells (Monds *et al*., 2006).

Such stress likely extends to other essential metabolic processes, triggering signaling pathways that activate mitigation strategies, including redox regulation when stress reaches the photosystems, as evidenced by the upregulation of DOMON-domain proteins and RBL-like serine proteases (Knopf & Adam, 2012; Eysholdt-Derzsó *et al*., 2023; Talukdar *et al*., 2024; Mukhtar *et al*., 2025). The increase in NPQ at later stages suggests that this stress extended to the photosystems, while the elevated qE indicates protective energy dissipation to protect the photosynthetic apparatus from redox-associated damage (Ruban, 2016). The likely involvement of the core carotenoid *t*-neoxanthin in this process is consistent with its proposed role as an alternative to the classical xanthophyll cycle in siphonous algae (Giossi *et al*., 2020, 2021). Moreover, the downregulation of hormone transduction pathways may represent reduced hormonal flexibility (Depuydt & Hardtke, 2011), while the upregulation of the *CCR4–NOT* complex may help maintain cellular homeostasis by facilitating selective transcript turnover during this later-stage stress (Collart, 2016; Zheng *et al*., 2023).

Overall, these findings suggest that in blue- and green-dominant light niches, *Derbesia* likely rapidly initiates acclimation to efficiently utilize these spectra for photosynthesis and sustained growth. Although some stress signals related to nutrient limitations emerged at the end of the experiment likely due to laboratory close culture systems, that is unlikely in nature. However, these stresses likely did not cause photosynthetic damage or morphological impairment. This is indicated by larger, rounded and reduced density of chloroplasts, the presence of all major photopigments and higher growth rates but likely represent compensatory responses that maintain optimal growth under these light conditions (Pyke & Leech, 1994).

### Long-wavelength light triggers a “low-light” survival strategy marked by morphological shifts and metabolic decline

In red (RL) and far-red (FL) conditions, *Derbesia* shows partial and strongly constrained photoacclimation, respectively. Under these spectra, the alga initially shows very low overall metabolic expressions, followed by limited activation of some of the core cellular processes at prolonged exposure; however, most pathways required for successful acclimation remain suppressed, particularly in FL, with only partial late recovery in RL. One possible explanation for the overall suppression of metabolism is that the 30 µmol photons m⁻² s⁻¹ of light in the RL and FL treatments delivered less overall energy usable by the alga than the BL, GL and WL treatments.

In FL and to a lesser extent in RL, changes in the light-dependent reactions were primarily associated with the regulation of LHC genes. The delayed induction of LHC genes likely indicates a later-emerging photoacclimation response aimed at expanding light-absorption capacity. This response is supported by elevated F_v_/F_m_ and α, along with increased accumulation of core antenna pigments, reflecting enlarged antenna size typical of low-light acclimation (Walters, 2005; Kirk, 2011; Dattolo *et al*., 2014; Zhen *et al*., 2019; Wu *et al*., 2022; Leschevin *et al*., 2024). The formation of narrow, densely packed chloroplasts with enlarged pyrenoids are also aimed at maximizing light capture and concentrating carbon fixation (Engel *et al*., 2015) as seen in low-light responses. However, despite antenna expansion, persistently high NPQ, qI, and qT_1_, and the absence of a marked increase in ΦPSII at later stages indicate inefficient utilization of the absorbed light energy as observed in higher plants (Ware *et al*., 2015; Taylor *et al*., 2019).

The high early expression of *MYCBP*, a transcriptional regulator associated with growth maintenance and stress signaling (Frank *et al*., 2001; Grandori *et al*., 2005; Liu *et al*., 2018), suggests that *Derbesia* initially restricted growth under FL, likely due to insufficient energy input to sustain normal development. Overexpression of *MYCBP* in *Arabidopsis* impairs vegetative growth and delays flowering (Kang *et al*., 2013), supporting the idea that elevated *MYCBP* activity reflects growth constraint under these light conditions.

Although Derbesia appeared to expand its light-harvesting capacity at later stages, the later activation of lipoxygenases and *SOD2*, consistent with enhanced lipid turnover, senescence, and ROS buffering (Hou *et al*., 2015; Gill *et al*., 2015; Palma *et al*., 2020; Singh *et al*., 2022) suggests active cellular stress during prolonged exposure, while growth remained restricted. Moreover, induction of linoleic acid metabolism, and glutathione metabolism, along with suppression of core growth-supporting processes including ATP metabolism, translation, DNA replication and cell-cycle, indicate stress mitigation rather than investment in growth, in line with the low growth compared to other light conditions. These observations are aligned with the reports linking far-red exposure to impaired cell division and physiological decline in algae (Sánchez-Saavedra *et al*., 1996; Zeebe *et al*., 1996; Xue *et al*., 2005). Additionally, strong induction of the chloroplast division gene *ftsZ*, accompanied by suppression of *ELIP* with loss of several accessory carotenoids resulted in large numbers of structurally remodelled chloroplasts under FL (Grimm *et al*., 1989; Bruno & Wetzel, 2004).

The high accumulation of core antenna pigments and the reduced Chl *a*/*b* ratio is likely the far-red acclimation strategy of *Derbesia.* Another bryopsidalean alga, *Ostreobium,* has also shown a reduced Chl *a*/*b* ratio along with the modification in Chl *a* molecule as a strategy for inhabiting far-red niches, suggesting this may be a lineage-wide far-red acclimation strategy within Bryopsidales (Koehne *et al*., 1999; Wilhelm & Jakob, 2006; Kotabová *et al*., 2014).

### Broad-spectrum white light results in peak performance

Full spectral white light (WL) showed photosynthetic adjustments largely similar to those observed under short-wavelength light conditions; however, the effective quantum yield of photosystem II, although initially similar to BL and GL, became highest under WL over time compared with all other light conditions. This likely reflects the complementary effects of short-and long-wavelength light, consistent with reports that supplementation of long wavelengths can enhance photochemical efficiency during photosynthesis (Emerson, 1957; Zhen *et al*., 2017, 2019).

Full spectral light strongly activated the core reaction-center components, ATP synthases, ATPases and ATP transporters during both early and prolonged exposure, indicating maintenance of photosystem integrity and sustained energy production and utilization throughout the experiment. Beyond photosynthesis, most genes associated with DNA replication, transcriptional regulation, and cytoskeletal organization also remained consistently highly expressed, indicating strong overall metabolic activity throughout the experiment. At the same time, due to the closed culture conditions and sustained high metabolic activity from the beginning, nutrient limitation was expected to emerge over time under WL, as reflected by the later-stage induction of the nutrient limitation marker *ApaH*-like phosphatases (Monds *et al*., 2006).

These consistently elevated metabolic responses suggest that *Derbesia* performs best in full-spectrum light conditions. This may reflect the evolutionary history of *Derbesia* in broad-spectrum sunlight in shallow coastal environments, where algae naturally experience fluctuating mixtures of wavelengths due to changes in water depth driven by tidal cycles, water movement, seasonal variation, and dissolved components (Kobara & Sears & Wilce, 1970; Schnetter *et al*., 1980; Chihara, 1981; Kirk, 2011; Strydom *et al*., 2017). Such long-term exposure may have shaped the alga’s metabolic systems to rely on broad spectral inputs for maintaining optimal physiological performance and successfully thriving within its natural coastal habitats.

## DECLARATION OF COMPETING INTEREST

The authors declare no competing financial interests or personal relationships that could have influenced the work presented in this study.

## ACKNOWLEDGEMENT

This work was supported by the Prime Minister’s Fellowship of the People’s Republic of Bangladesh to RH; a fellowship from the Fundação para a Ciência e a Tecnologia to HV (https://doi.org/10.54499/2023.06155.CEECIND/CP2845/CT0004); and Gorden Betty Moore foundation, and Mary Lugton fellowship by the University of Melbourne to PD. RH also acknowledges support from the Dr David Lachlan Hay Memorial Postgraduate Writing-Up Award by the University of Melbourne. We also thank the HMSTrust Analytical Laboratory at Monash University for their support with pigment analysis via HPLC and for access to their facilities. We are also grateful to the members of the Verbruggen Laboratory for their insightful suggestions regarding data interpretation, and to Dr. Md Abdullah Al Kamran Khan for his assistance during sampling.

## Notes

### Competing Interest Statement

The authors have declared no competing interest.

