## Supplemental file for "Thriving to surviving: light wavelength modulates photoacclimation response in the siphonous green alga *Derbesia*"

**Method S1. Photosynthesis measurements using chlorophyll a fluorescence**

Photosynthetic activity was assessed using an Imaging-PAM fluorometer (Maxi version; Heinz Walz GmbH, Germany) through light induction and recovery curves, as well as rapid light curves. Macroalgal samples were dark acclimated for 20 minutes before measurements, and chlorophyll fluorescence parameters were obtained using routine saturating pulse analyses throughout the sampling days. Fluorescence parameters were extracted from digital images using ImagingWin software by selecting areas of interest. The saturating light pulse analyses provided key PSII activity parameters. The maximum quantum yield of PSII ($F_v/F_m$) was calculated as ($F_m$ - $F_0$) / $F_m$, where $F_m$ and $F_0$ represent the maximum and minimum fluorescence of dark-acclimated samples. Under continuous illumination, the effective PSII quantum yield was determined as ($F_m'$ - F) / $F_m'$, where $F_m'$ and F correspond to the maximum and transient fluorescence, respectively. Non-photochemical quenching (NPQ) was calculated as $F_m$/ $F_m'$ - 1, where $F_m'$ represents the last maximum quantum yield under actinic light conditions in the induction curve. The fraction of PSII reaction centers in the open state was estimated using the qP parameter, defined as {( $F_m'$ - F) / ($F_m'$ - $F_0'$)} × ($F_0'$ / F), where $F_0'$ is a theoretical estimation of the minimum fluorescence in light-acclimated samples, calculated as described by Oxborough and Baker (1997).

Rapid light curves (RLCs) were measured without prior dark acclimation. Samples were taken directly from their culture conditions and exposed to a series of actinic light intensities, with each light step lasting 60 seconds. A saturating light pulse was applied at the end of each step to estimate the relative electron transport rate (rETR) of PSII, calculated as 0.42 × ($F_m'$ - F) / $F_m'$) × PPFD, where 0.42 represents the estimated fraction of incident photons absorbed by PSII. Light response curves were fitted to the model of Platt *et al.* (1980) to estimate two key parameters: the initial slope (α) of the ETR versus irradiance curve, and the maximum relative ETR (rETRmax). Curve fitting was performed iteratively using MS Excel Solver, and the fits were consistently considered reliable (r > 0.98).

All measurements were conducted in a dedicated imaging room while maintaining a constant temperature of 26 ± 0.5°C throughout the iPAM protocol. The measuring beam and actinic light of the Imaging-PAM were both blue, and the general settings for Imaging-PAM were ML = 3, Frequency = 2, and Actinic light step= 5 (81 µmol $m^2$ s). Dark incubation was achieved by wrapping cultures in aluminium foil while keeping them in their original growth conditions. To ensure consistency, all fluorescence measurements were performed at the same time point, beginning at 10 AM, four hours after the onset of light.

**Table S1.** List of photosynthetic parameters estimated based on chl *a* fluorescence induction and relaxation kinetics, rapid light curve (RLC) fitting and de-epoxidation state (DES).

| Photosynthetic parameters | Formulae | References |
|---|---|---|
| Maximum quantum yield of PSII, $F_v/F_m$ | $\frac{F_m - F_0}{F_m}$ | (Klughammer and Schreiber 2008) |
| Effective quantum yield, ΦPSII | $\frac{F_m' - F_0'}{F_m'}$ | (Klughammer and Schreiber 2008) |
| Co-efficient of photochemical quenching based on puddle model, qP | $\frac{F_m' - F}{F_m' - F_0'}$ | (Oxborough and Baker 1997) |
| Non-photochemical quenching Stern volmer, NPQ | $\frac{F_m}{F_m'} - 1$ | (Klughammer and Schreiber 2008) |
| Electron transport rate, ETR | $\frac{F_v}{F_m} * 0.84 * 0.5 * PAR$ | (Schreiber 2004) |
| Coefficient of non-photochemical quenching, qN | $\frac{F_m - F_m'}{F_m - F_0'}$ | (Schreiber and Klughammer 2008) |
| Quantum yield of regulated energy dissipation, Y(NPQ) | $1 - \frac{(F_m' - F)}{F_m} - \frac{1}{(NPQ + 1 + qL\left(\frac{F_m}{F_0}\right) - 1)}$ | (Kramer et al. 2004) |
| Quantum yield of non-regulated energy dissipation, Y(NO) | $\frac{1}{(NPQ + 1 + qL\left(\frac{F_m}{F_0} - 1\right)}$ | (Kramer et al. 2004) |
| Minimum saturation irradiance, Ek | $E_k = \frac{ETR_{max}}{\alpha}$ | (Ralph and Gademann 2005) |

| | | |
|---|---|---|
| Fast relaxing NPQ, qE type | $\left(\frac{F_m - F_m'5}{F_m'5}\right) - \left(\frac{F_m - maxF_m'6\text{-}20}{maxF_m'6\text{-}20}\right)$ | (Aihara et al. 2016, Herdean et al. 2023) |
| Slow relaxing NPQ, **qI** type | $\left(\frac{F_m - maxF_m'6\text{-}20}{maxF_m'6\text{-}20}\right)$ | (Herdean et al. 2023) |
| State transition I, $qT_1$ type | $\left(\frac{F_m in\ light - maxF_m'6\text{-}20}{maxF_m'6\text{-}20}\right) - \left(\frac{F_m in\ dark - maxF_m'6\text{-}20}{maxF_m'6\text{-}20}\right)$ | (Herdean et al. 2023) |

### Cited references of this table

Aihara Y., Takahashi, S. & Minagawa, J. 2016. Heat induction of cyclic electron flow around photosystem I in the symbiotic dinoflagellate *Symbiodinium*. *Plant Physiology* 171:522-529.

Herdean A., Hall, C., Hughes, D. J., Kuzhiumparambil, U., Diocaretz, B. C. & Ralph, P. J. 2023. Temperature mapping of non-photochemical quenching in Chlorella vulgaris. *Photosynthesis Research* 155:191-202.

Klughammer C. & Schreiber, U. 2008. Complementary PSII quantum yields calculated from simple fluorescence parameters measured by PAM fluorometry and the Saturation Pulse method. *PAM Application Notes* 1:201-247.

Kramer D. M., Johnson, G., Kiirats, O. & Edwards, G. E. 2004. New fluorescence parameters for the determination of QA redox state and excitation energy fluxes. *Photosynthesis Research* 79:209-218.

Oxborough K. & Baker, N. R. 1997. Resolving chlorophyll a fluorescence images of photosynthetic efficiency into photochemical and non-photochemical components–calculation of qP and $F_v/F_m$; without measuring $F_o$. *Photosynthesis Research* 54:135-142.

Ralph P. J. & Gademann, R. 2005. Rapid light curves: a powerful tool to assess photosynthetic activity. *Aquatic Botany* 82:222-237.

Schreiber U. 2004. Pulse-Amplitude-Modulation (PAM) Fluorometry and Saturation Pulse Method: An Overview. *Chlorophyll a Fluorescence: A Signature of Photosynthesis*. Papageorgiou, G. C. and Govindjee [eds]. Springer Netherlands, Dordrecht, pp. 279-319.

Schreiber U. & Klughammer, C. 2008. Non-photochemical fluorescence quenching and quantum yields in PSI and PSII: analysis of heat-induced limitations using Maxi-Imaging-PAM and Dual-PAM-100. *PAM Application Notes* 1:15-18.

**Table S2.** Absorption spectra and retention time of the identified pigments in this study following the previous works of Giossi *et al.*, (2021); Mendes *et al.*, (2007). Note- the observed retention time and absorption maxima shows different from their reference value due to the different solvent gradient used.

| Pigments | Abbreviation | Retention time (min) | Ref Retention Time (min) | absorption maxima (nm) | Ref absorption maxima (nm) |
|---|---|---|---|---|---|
| Siphonoxanthin | Sip | 3.99 | 10.41 | 454 | 448 |
| All-trans-neoxanthin | *t*-Neo | 4.65 | 11.38 | 420, 443, 472 | 418, 441, 470 |
| 9'-cis-Neoxanthin | *c*-Neo | 4.93 | 12 | 416, 439, 467 | 413, 437, 466 |
| Violaxanthin | Viola | 5.18 | 13 | 419, 442, 472 | 416, 441, 471 |
| Zeaxanthin | Zea | 6.9 | 18.71 | 455, 480 | 430, 454, 481 |
| Lutein | Lut | 7.07 | 18.24 | 448, 476 | 425, 447, 475 |
| Siphonaxanthin dodecenoate | Sip-d | 8.00 | 18.53 | 457 | 455 |
| Chlorophyll-b | Chl-b | 8.52 | 22.85 | 460, 646 | 458, 595, 645 |
| Chlorophyll-a | Chl-a | 8.92 | 24.22 | 430, 665 | 430, 617, 663 |
| β,ε-carotene | ε-Car | 9.78 | 28.14 | 415, 442, 471 | 418, 441, 471 |
| α-Carotene | α-Car | 9.90 | 28.33 | 447, 475 | 447, 476 |
| Unknown carotenoid | Un-car | 5.4 | – | 447 | – |
| Derivative of neoxanthin | D-Neo | 3.5 | – | 419;444;471 | – |

**Method S2. Methods for statistical analysis for iPAM data**

Statistical analyses were performed using R (version 4.4.0; R Core Team, 2024). Before analysis, data were screened for technical artifacts; values of zero resulting from failed data acquisition or extraction were treated as missing data (NA) to prevent bias in variance estimation and mean calculation. To evaluate the physiological responses across treatments and time points, a linear model was constructed for each chlorophyll *a* fluorescence parameter. The models included Treatment (5 levels: Control, Blue, Green, Red, and Far-red) and Sampling Day (2 levels: Day 4 and Day 10) as fixed effects, including their interaction term (Treatment\Day).

Due to the missing replicates in some cases, Estimated Marginal Means (EMMs)—also known as least-squares means—were calculated using the emmeans package (Lenth, 2023). This approach provides unbiased mean estimates by utilizing the pooled variance of the entire dataset. Two specific sets of planned comparisons (simple effects) were conducted:

(i) Temporal Effects: Pairwise comparisons between Day 4 and Day 10 were performed within each treatment to assess physiological shifts over the experimental period.

(ii) Treatment Effects: Differences among treatment conditions were evaluated independently at each sampling time point (Day 4 and Day 10) to identify specific light-quality responses.

All post-hoc pairwise comparisons were adjusted for multiple testing using Tukey's Honestly Significant Difference (HSD) method to control the family-wise error rate. Results were considered statistically significant at $p < 0.05$. Data are presented as the mean standard error of the mean (SEM).

**Table S3:** Significance level from the statistical testing of iPAM parameters with indicating asterisk for significant and "ns" for non-significant values. Here, D means Sampling day, *p* means significance level, *p* value; "-" means fails calculation due to missing data.

| Parameter | *P* (Treatment) | *P* (across treatm_D4) | *P* (across treatm_D10) | *P* (D4vD10_Control) | *P* (D4vD10_Blue) | *p* (D4vD10_Green) | *p* (D4vD10_Red) | *p* (D4vD10_Far-red) |
|---|---|---|---|---|---|---|---|---|
| $F_v/F_m$ | *** | *** | *** | ns | ns | *** | *** | ns |
| ΦPSII | ns | ** | ns | * | ns | - | ** | ns |
| 1-qP | ns | ** | ns | - | - | ns | * | ns |
| NPQ | ** | * | ns | ns | - | ns | ns | ns |
| Y(NPQ) | ns | * | ns | ns | - | ns | ns | ** |
| Y(NO) | *** | ** | * | ns | - | ns | ns | * |
| ql | *** | ns | ** | ns | - | ns | ns | ns |
| qE | ns | ns | ns | - | ns | - | - | * |

| | | | | | | | | |
|---|---|---|---|---|---|---|---|---|
| $qT_1$ | ** | ns | * | ns | - | ns | ns | ns |
| Alpha | *** | *** | *** | ns | ** | ** | ns | ns |
| $E_k$ | *** | *** | *** | ** | ns | *** | ** | ** |

**Table S4:** The numeric *p* value of the statistical testing of iPAM parameters across the treatments and sampling days

| Parameter | *p* (Treatment) | *p* (across treatm_D4) | *p* (across treatm_D10) | *p* (D4vD10_Control) | *p* (D4vD10_Blue) | *p* (D4vD10_Green) | *p* (D4vD10_Red) | *p* (D4vD0_Far-red) |
|---|---|---|---|---|---|---|---|---|
| $F_v/F_m$ | 2.72E-12 | 7.96E-06 | 3.01E-07 | 0.207043 | 0.151755 | 0.000198 | 0.000689 | 0.26088 |
| ΦPSII | 0.182192 | 0.001472 | 0.075184 | 0.046265 | 0.239715 | - | 0.002582 | 0.146465 |
| 1-qP | 0.497545 | 0.001949 | 0.326274 | - | - | 0.170499 | 0.013997 | 0.063591 |
| NPQ | 0.001266 | 0.045995 | 0.052634 | 0.986545 | - | 0.576971 | 0.362344 | 0.124601 |
| Y(NPQ) | 0.595673 | 0.032625 | 0.68079 | 0.402795 | - | 0.094733 | 0.327043 | 0.009963 |
| Y(NO) | 7.55E-05 | 0.007856 | 0.014301 | 0.95482 | - | 0.260888 | 0.905576 | 0.014803 |
| ql | 0.00041 | 0.1619 | 0.009124 | 0.92043 | - | 0.567571 | 0.170952 | 0.09948 |
| qE | 0.820328 | 0.789514 | 0.645762 | - | 0.854216 | - | - | 0.025789 |
| $qT_1$ | 0.001554 | 0.1619 | 0.024663 | 0.92043 | - | 0.390008 | 0.150053 | 0.09948 |
| Alpha | 4.25E-13 | 6.79E-07 | 6.27E-07 | 0.078698 | 0.004891 | 0.004988 | 0.575832 | 0.066612 |
| $E_k$ | 1.66E-09 | 0.000129 | 1.60E-08 | 0.006788 | 0.05482 | 7.68E-05 | 0.004303 | 0.008062 |

**Table 5.** The *p* value of the statistical testing of growth rates, the parameters of chloroplast morphology, and the photopigments across the treatments. Here, 1= White light, 2 = Blue light, 3 = Green light, 4 = Red light, 5 = Far-red light) at *P* <5%.

| Group | Sum of Square | Mean Square | F Statistic | P-value |
|---|---|---|---|---|
| Growth rate | 604.5299 | 151.1325 | 391.3756 | 0 |
| 1v2 | | | | 0.9078 |
| 1v3 | | | | 0.4966 |
| 1v4 | | | | 2.203e-10 |
| 1v5 | | | | 5.11e-13 |
| 2v3 | | | | 0.9365 |
| 2v4 | | | | 7.139e-11 |
| 2v5 | | | | 5.11e-13 |
| 3v4 | | | | 2.769e-11 |
| 3v5 | | | | 5.11e-13 |
| 4v5 | | | | 1.679e-12 |
| Cp density | 0.08057 | 0.02014 | 60.0822 | 0 |
| 1v2 | | | | 0.9935 |
| 1v3 | | | | 0.9733 |
| 1v4 | | | | 0.000006336 |
| 1v5 | | | | 1.315e-10 |
| 2v3 | | | | 0.9997 |
| 2v4 | | | | 0.000001747 |
| 2v5 | | | | 1.315e-10 |
| 3v4 | | | | 0.000001149 |
| 3v5 | | | | 1.315e-10 |
| 4v5 | | | | 9.751e-10 |
| Roundness | 7.7505 | 1.9376 | 242.5156 | 0 |
| 1v2 | | | | 0.9998 |
| 1v3 | | | | 1 |

| | | | | |
|---|---|---|---|---|
| 1v4 | | | | 7.786e-8 |
| 1v5 | | | | 1.315e-10 |
| 2v3 | | | | 0.9999 |
| 2v4 | | | | 4.67e-7 |
| 2v5 | | | | 1.315e-10 |
| 3v4 | | | | 4.034e-7 |
| 3v5 | | | | 1.315e-10 |
| 4v5 | | | | 1.315e-10 |
| alpha_carotene | 0.01588 | 0.003971 | 18.9514 | 0.0001165 |
| 1v2 | | | | 0.7351 |
| 1v3 | | | | 1 |
| 1v4 | | | | 0.0007662 |
| 1v5 | | | | 0.8199 |
| 2v3 | | | | 0.7615 |
| 2v4 | | | | 0.0001723 |
| 2v5 | | | | 0.9998 |
| 3v4 | | | | 0.000719 |
| 3v5 | | | | 0.8428 |
| 4v5 | | | | 0.000208 |
| Chlorophyll *b* | 0.2981 | 0.07453 | 89.7034 | 8.594e-8 |
| 1v2 | | | | 0.8528 |
| 1v3 | | | | 0.9875 |
| 1v4 | | | | 0.00001467 |
| 1v5 | | | | 0.00003001 |
| 2v3 | | | | 0.5995 |
| 2v4 | | | | 0.000006067 |
| 2v5 | | | | 0.00008343 |
| 3v4 | | | | 0.0000231 |
| 3v5 | | | | 0.00001886 |

| | | | | |
|---|---|---|---|---|
| 4v5 | | | | 3.211e-8 |
| Siph-d | 0.2307 | 0.05767 | 87.2328 | 9.838e-8 |
| 1v2 | | | | 0.00001129 |
| 1v3 | | | | 0.00001468 |
| 1v4 | | | | 0.00001328 |
| 1v5 | | | | 0.02467 |
| 2v3 | | | | 0.9983 |
| 2v4 | | | | 0.9997 |
| 2v5 | | | | 5.907e-7 |
| 3v4 | | | | 1 |
| 3v5 | | | | 7.194e-7 |
| 4v5 | | | | 6.671e-7 |
| Lut | 0.0001222 | 0.00004073 | 5.0042 | 0.03052 |
| 1v2 | | | | 0.06067 |
| 1v3 | | | | 0.02968 |
| 1v4 | | | | 0.1551 |
| 2v3 | | | | 0.9542 |
| 2v4 | | | | 0.9061 |
| 3v4 | | | | 0.6553 |
| Zeax | 0.003913 | 0.001304 | 2.4695 | 0.1364 |
| Un-car | 0.0005986 | 0.0002993 | 20.2958 | 0.002136 |
| 1v2 | | | | 0.003243 |
| 1v3 | | | | 0.004017 |
| 2v3 | | | | 0.9707 |
| Viol | 0.01349 | 0.003372 | 4.3017 | 0.02792 |
| 1v2 | | | | 0.9944 |
| 1v3 | | | | 0.5205 |
| 1v4 | | | | 0.2035 |
| 1v5 | | | | 0.9824 |

| | | | | |
|---|---|---|---|---|
| 2v3 | | | | 0.7373 |
| 2v4 | | | | 0.115 |
| 2v5 | | | | 0.9999 |
| 3v4 | | | | 0.01735 |
| 3v5 | | | | 0.8089 |
| 4v5 | | | | 0.09314 |
| *c*-neo | 0.5656 | 0.1414 | 89.0479 | 8.905e-8 |
| 1v2 | | | | 0.06661 |
| 1v3 | | | | 0.1687 |
| 1v4 | | | | 0.1807 |
| 1v5 | | | | 0.000001358 |
| 2v3 | | | | 0.9693 |
| 2v4 | | | | 0.9602 |
| 2v5 | | | | 1.755e-7 |
| 3v4 | | | | 1 |
| 3v5 | | | | 2.527e-7 |
| 4v5 | | | | 2.601e-7 |
| *t*-neo | 0.02563 | 0.006408 | 39.6962 | 0.000004164 |
| 1v2 | | | | 0.4536 |
| 1v3 | | | | 0.2315 |
| 1v4 | | | | 0.000401 |
| 1v5 | | | | 0.000447 |
| 2v3 | | | | 0.9824 |
| 2v4 | | | | 0.000056 |
| 2v5 | | | | 0.000061 |
| 3v4 | | | | 0.000032 |
| 3v5 | | | | 0.000035 |
| 4v5 | | | | 1 |
| Siph | 1.33 | 0.3325 | 117.8334 | 2.282e-8 |

| | | | | |
|---|---|---|---|---|
| 1v2 | | | | 0.03458 |
| 1v3 | | | | 0.05341 |
| 1v4 | | | | 0.1695 |
| 1v5 | | | | 3.667e-7 |
| 2v3 | | | | 0.9984 |
| 2v4 | | | | 0.8311 |
| 2v5 | | | | 4.884e-8 |
| 3v4 | | | | 0.9362 |
| 3v5 | | | | 5.684e-8 |
| 4v5 | | | | 8.595e-8 |
| D-neo | | | | 0.005118 |
| 1v2 | | | | 0.01575 |
| 1v3 | 0.00001851 | 0.000009254 | 14.4078 | 0.005468 |
| 2v3 | | | | 0.593 |

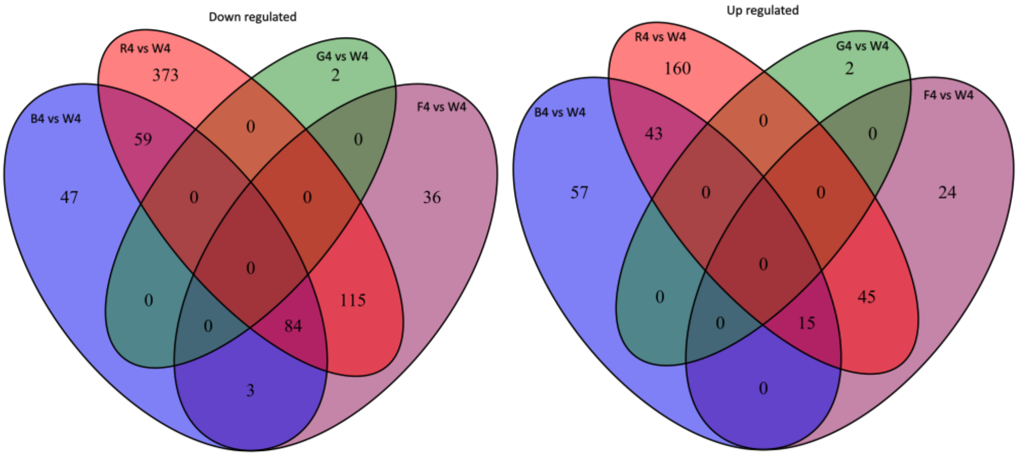

**Figure S1:** Venn diagram of expressed genes of treatment vs White light at day4.

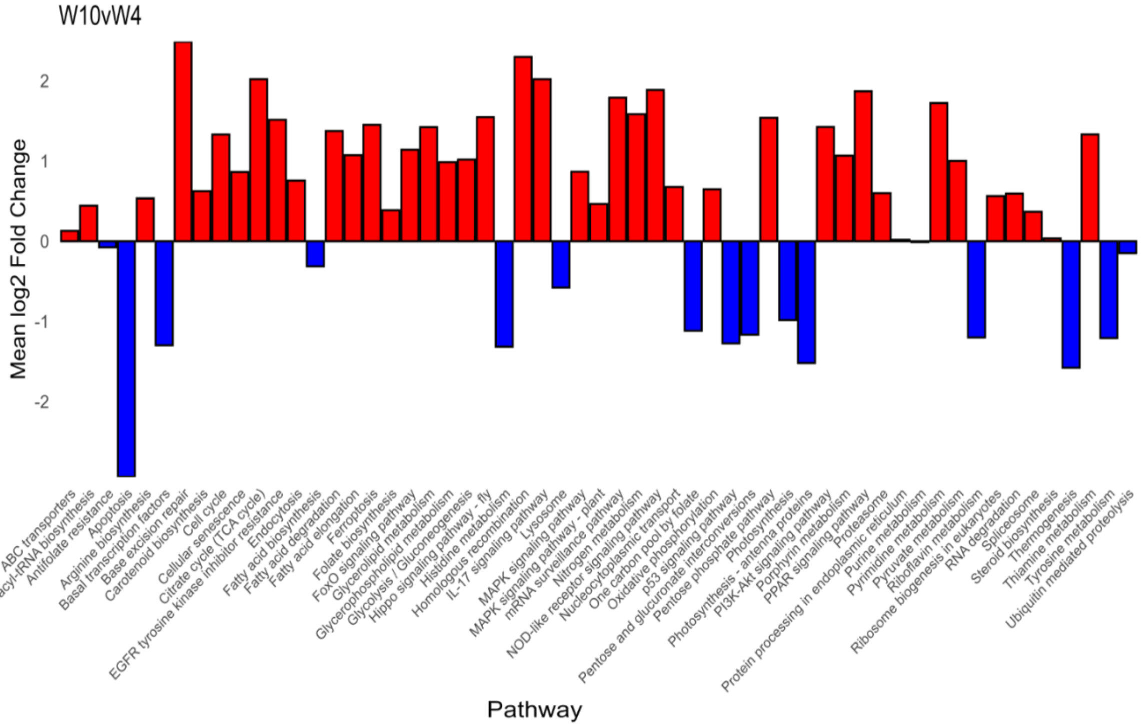

**Figure S2:** KEGG pathways change within White light at its Day10 vs. Day4

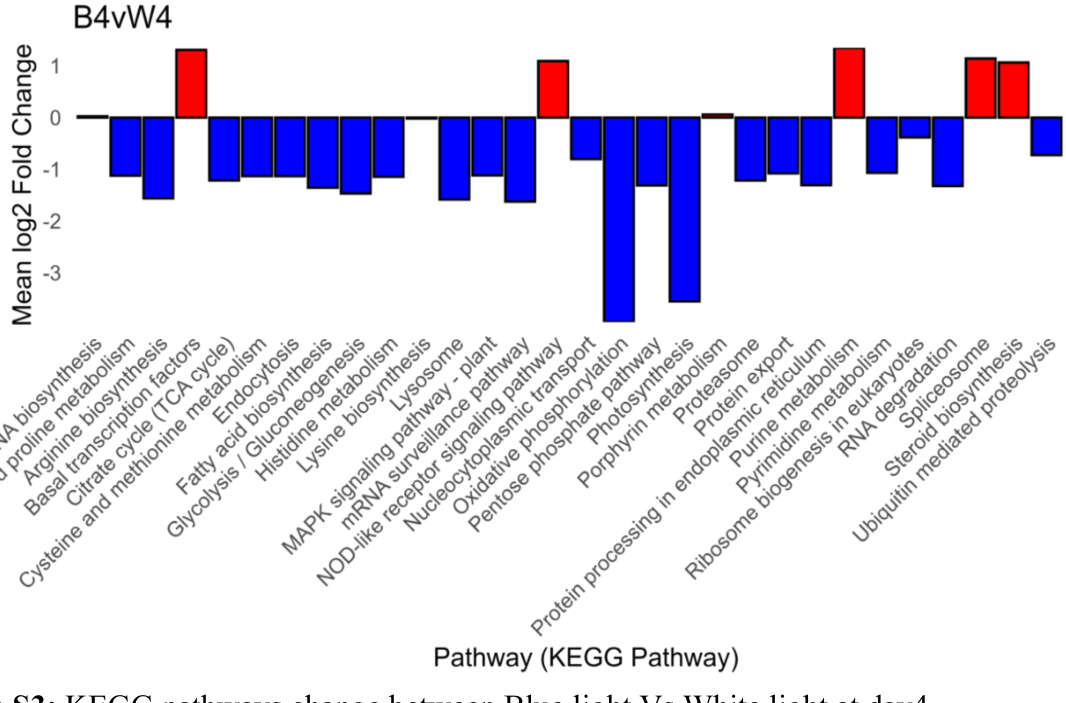

**Figure S3:** KEGG pathways change between Blue light Vs White light at day4.

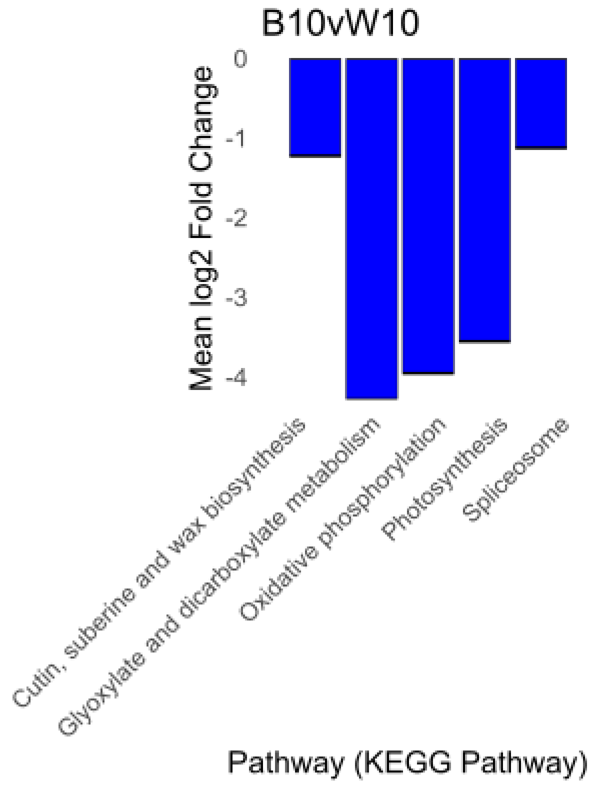

**Figure S4:** KEGG pathways change between Blue light Vs White light at day10.

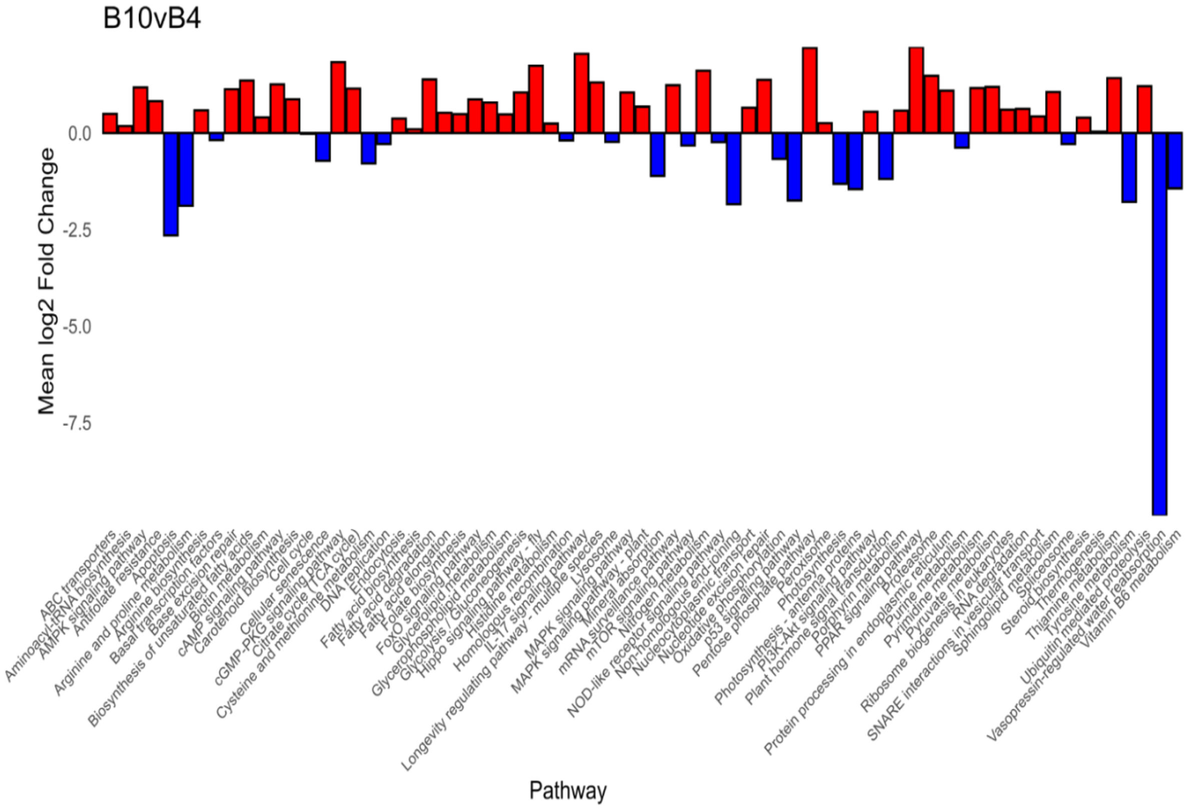

**Figure S5:** KEGG pathways change within Blue light at day 10 vs. day 4.

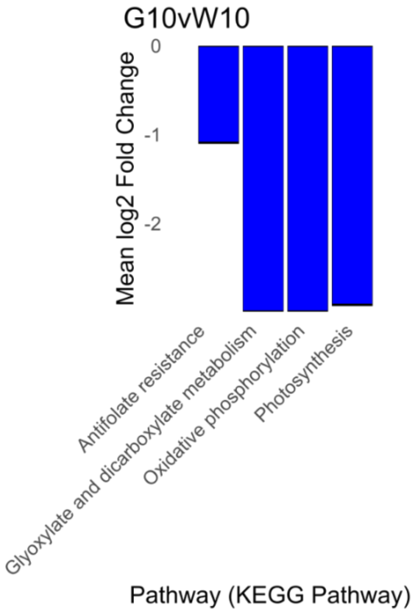

**Figure S6:** KEGG pathways change between Green light vs. White light of day 10.

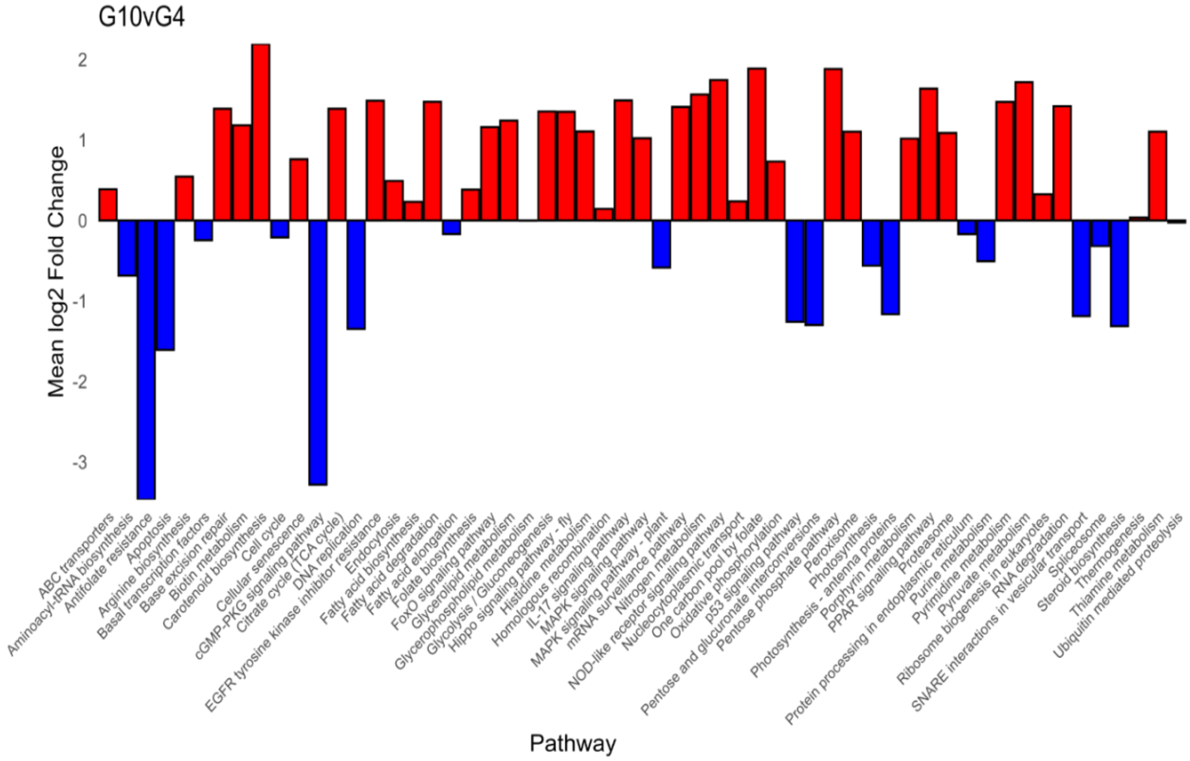

**Figure S7:** KEGG pathways change within Green light at day 10 vs. day 4.

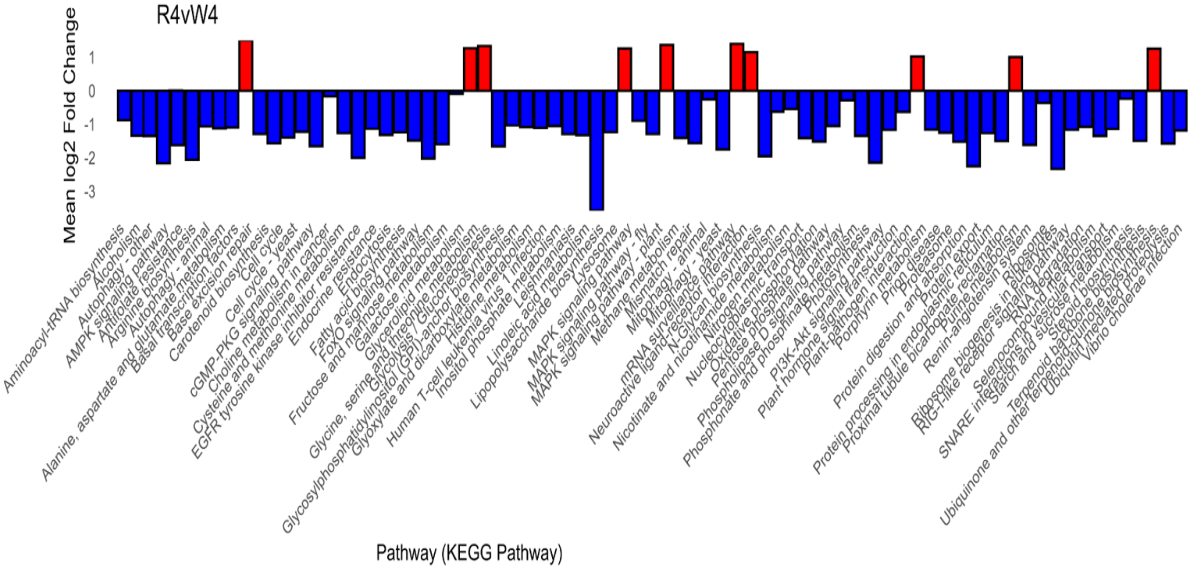

**Figure S8:** KEGG pathways change between Red light vs. White light of day 4.

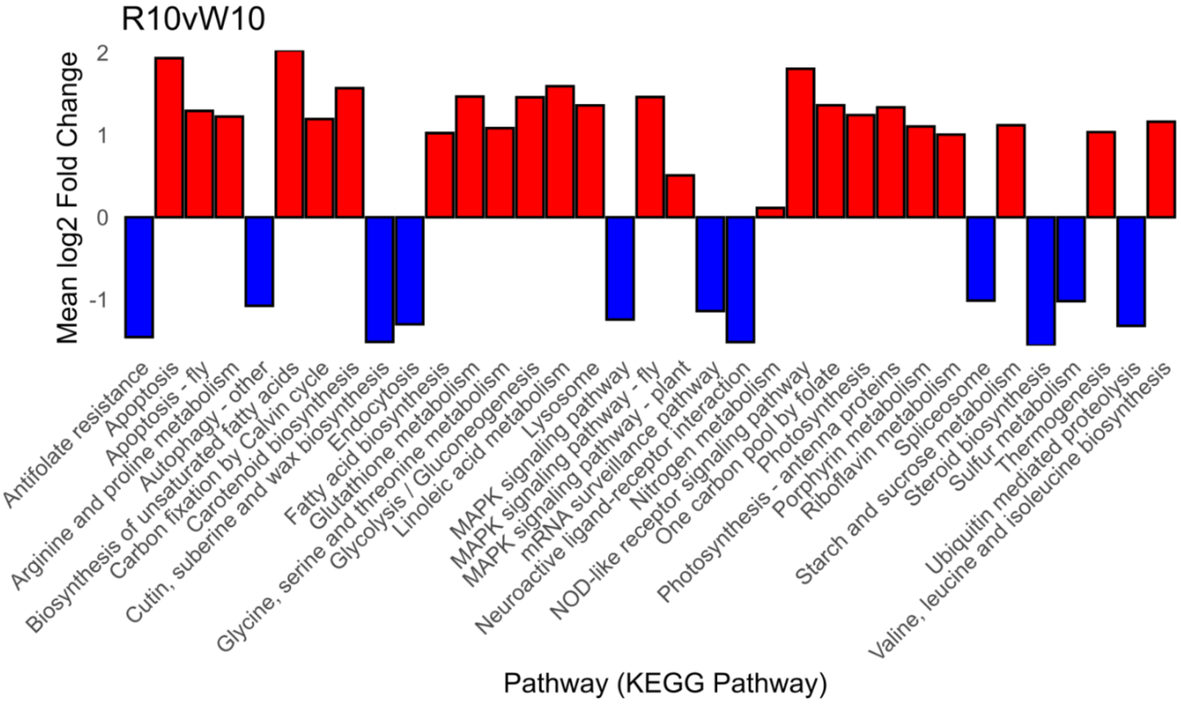

**Figure S9:** KEGG pathways change between Red light vs. White light of day 10.

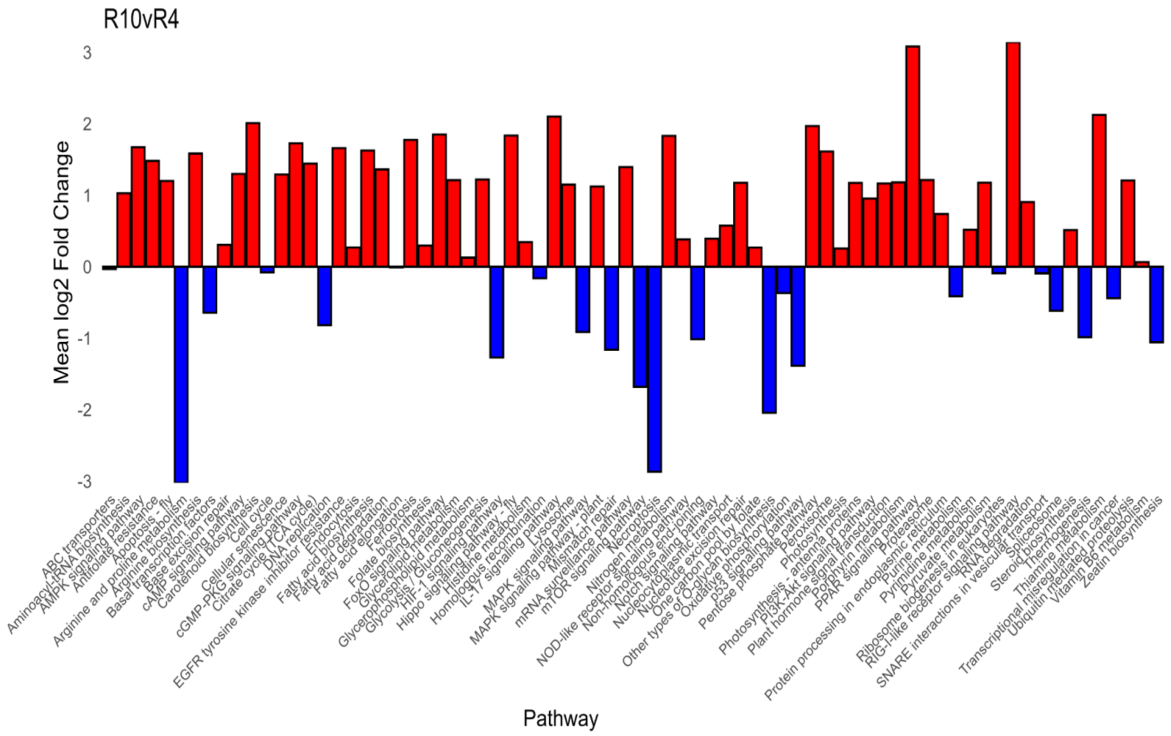

**Figure S10:** KEGG pathways change within Red light at day 10 vs. day 4.

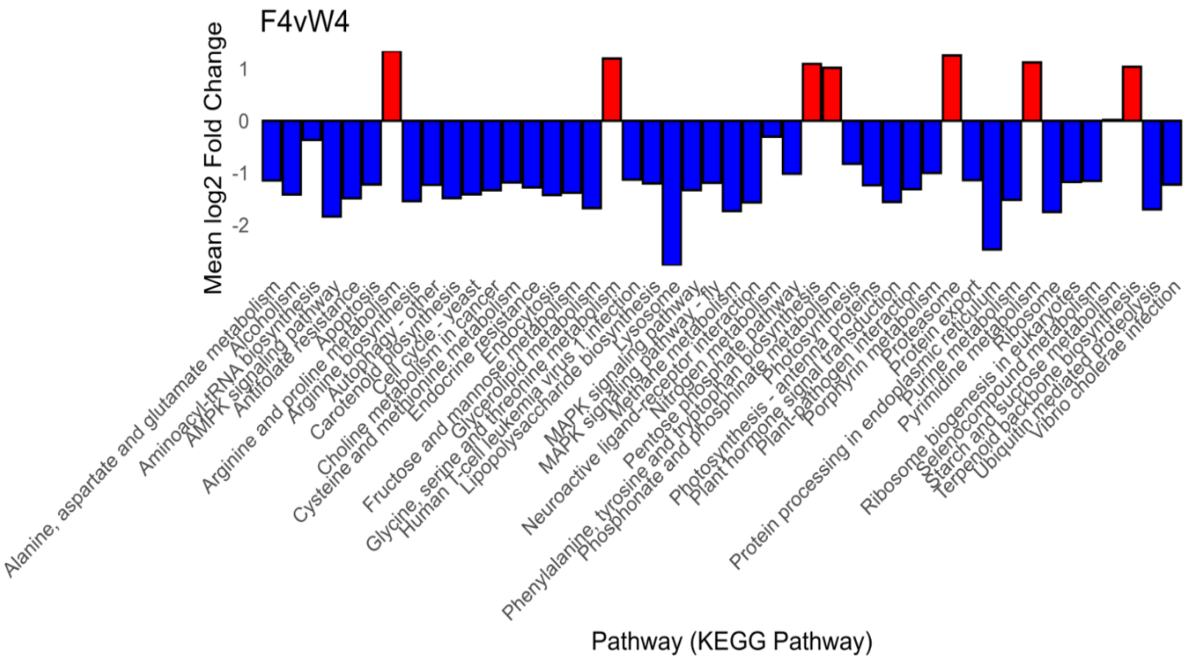

**Figure S11:** KEGG pathways change between Far-red light vs. White light of day 4.

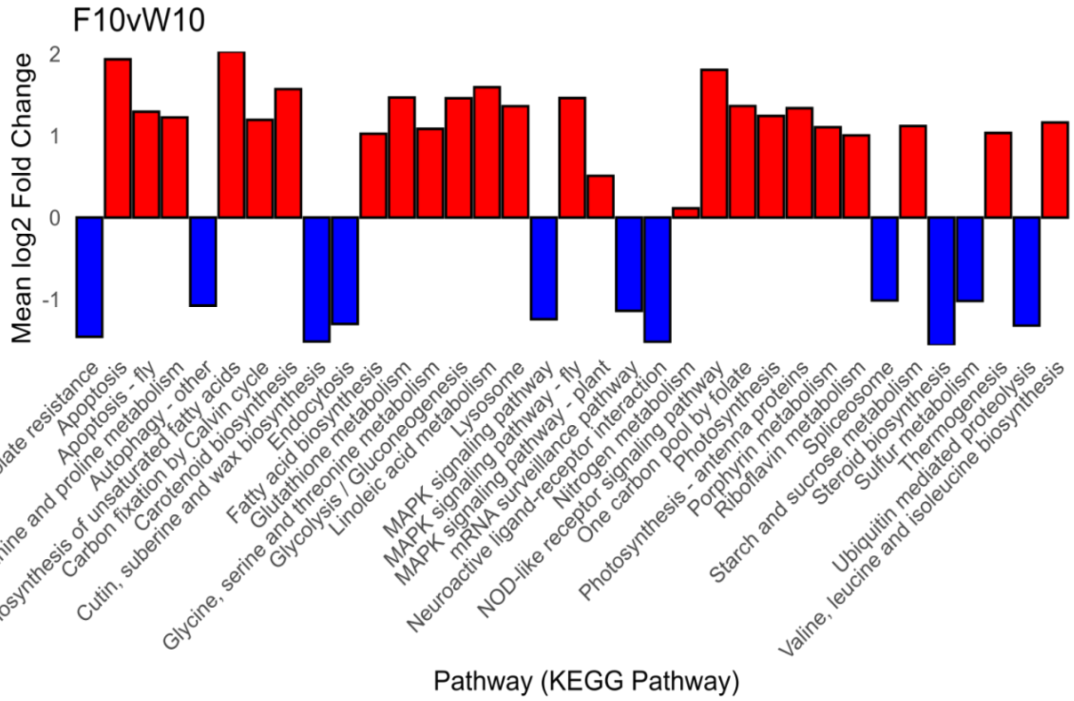

**Figure S12:** KEGG pathways change between Far-red light vs. White light of day 10.

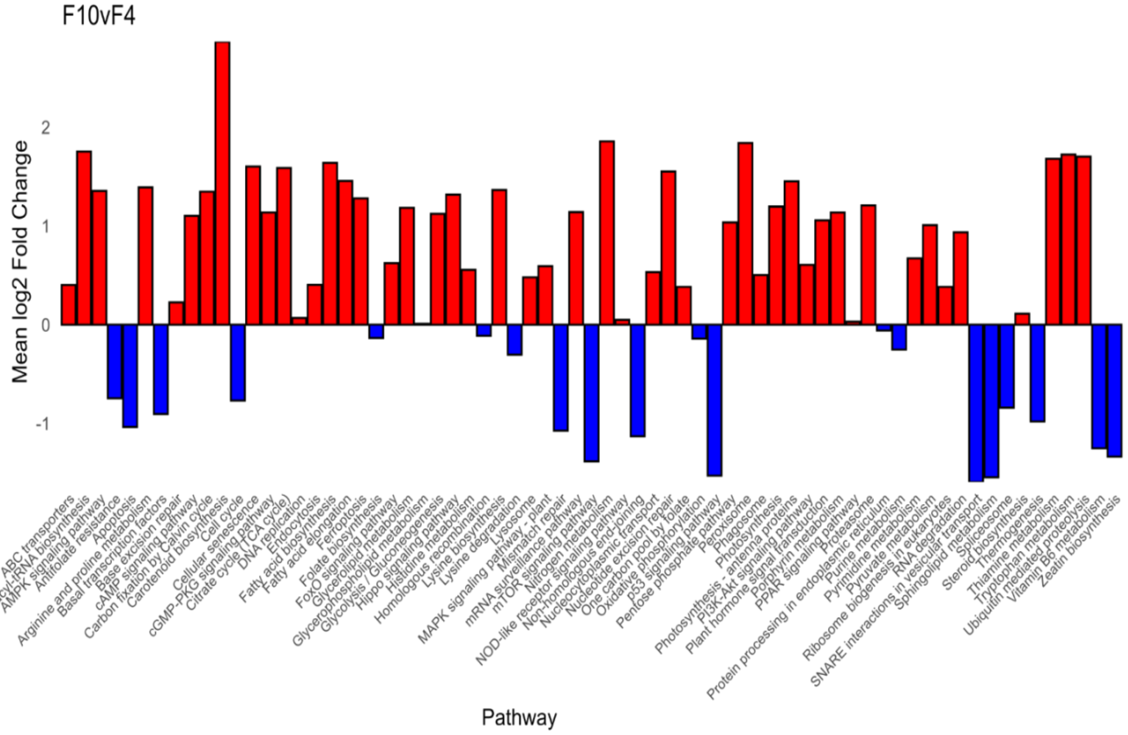

**Figure S13:** KEGG pathways change within Far-red light at day 10 vs. day 4.

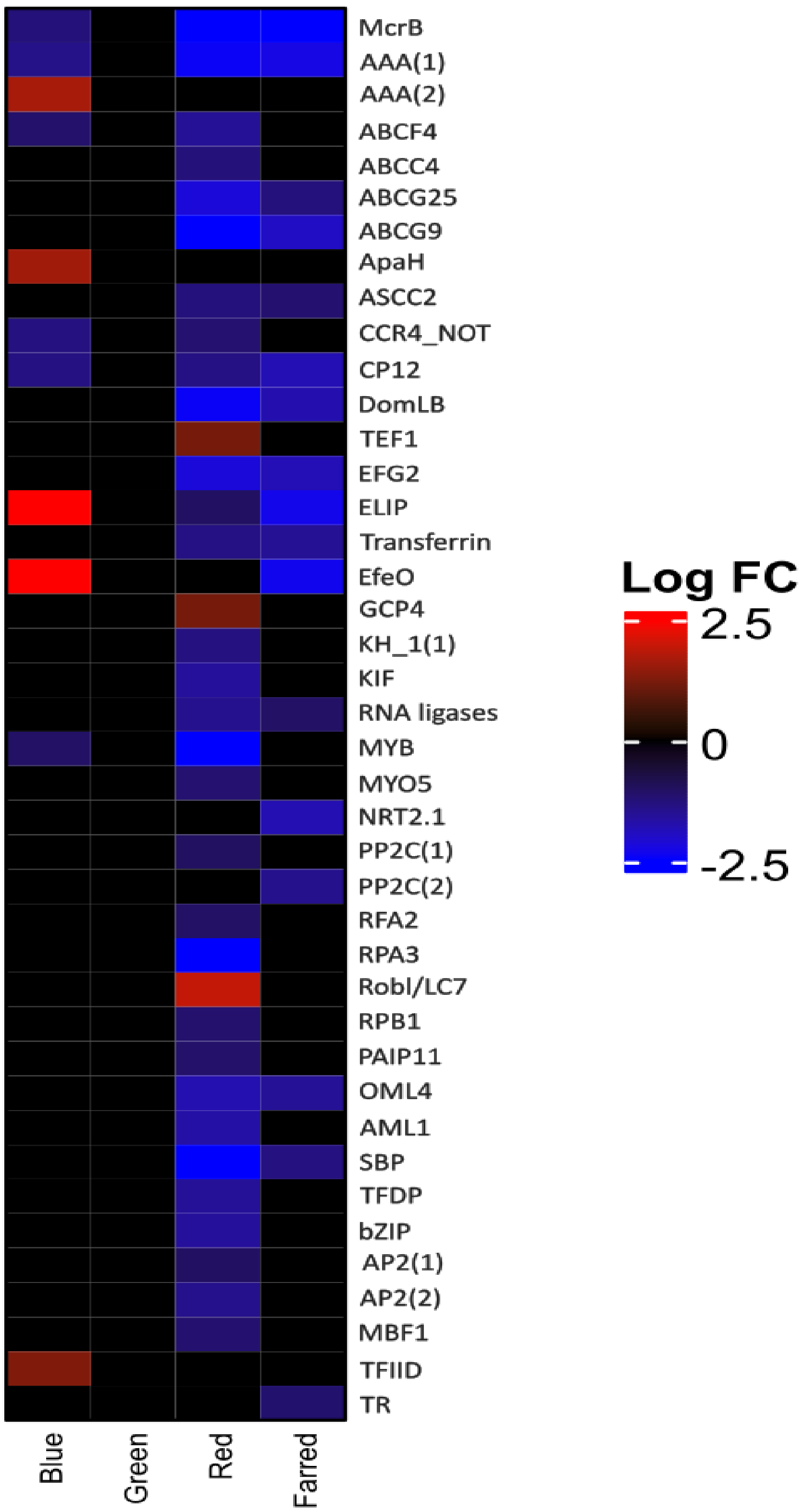

**Figure S14.** Heatmap showing the common DEGs identified in treatment vs white light at Day 4, but commonly showing DEGs when comparing Day 10 vs Day 4.

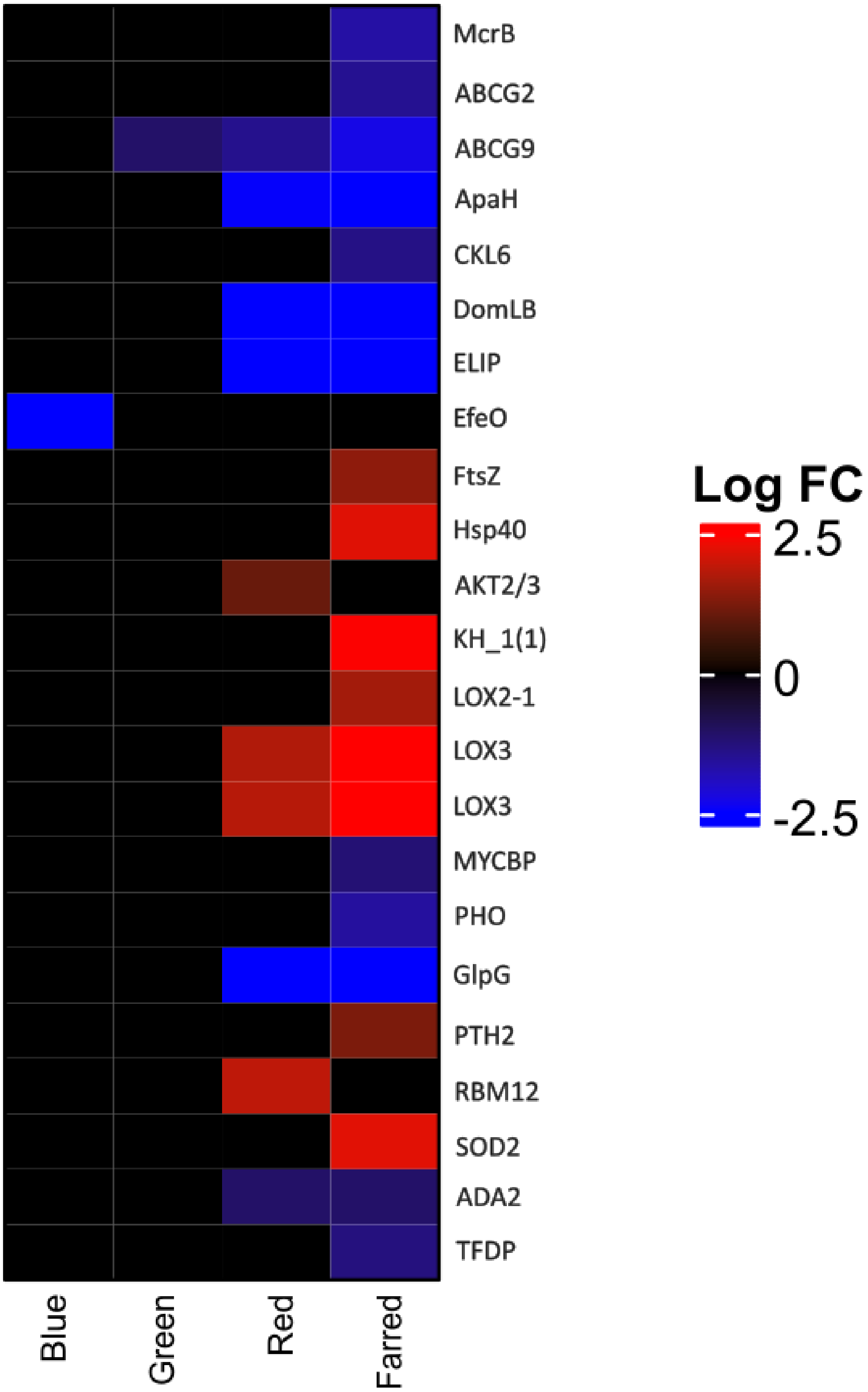

**Figure S15.** Heatmap showing the common DEGs identified in treatment vs white light at Day 10, but commonly showing DEGs when comparing Day 10 vs Day 4.

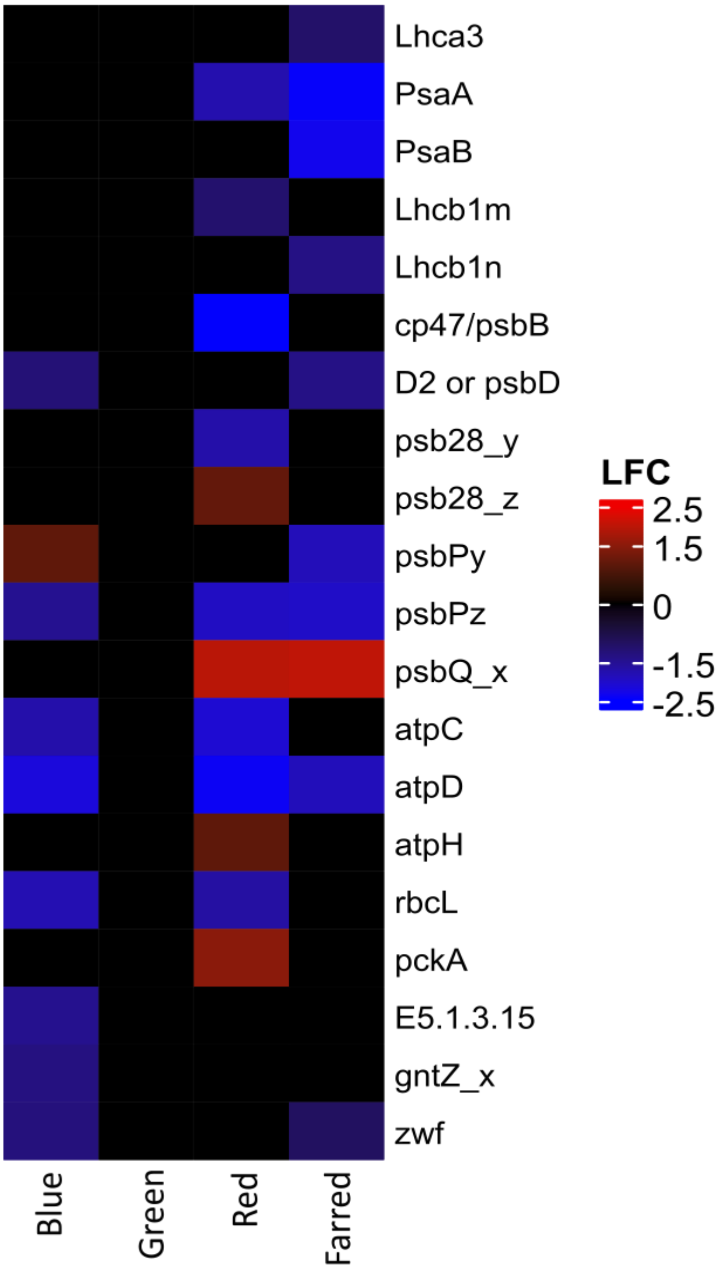

**Figure S16.** Heatmap of photosynthesis, and central carbohydrate metabolism genes of treatment vs White light at day4

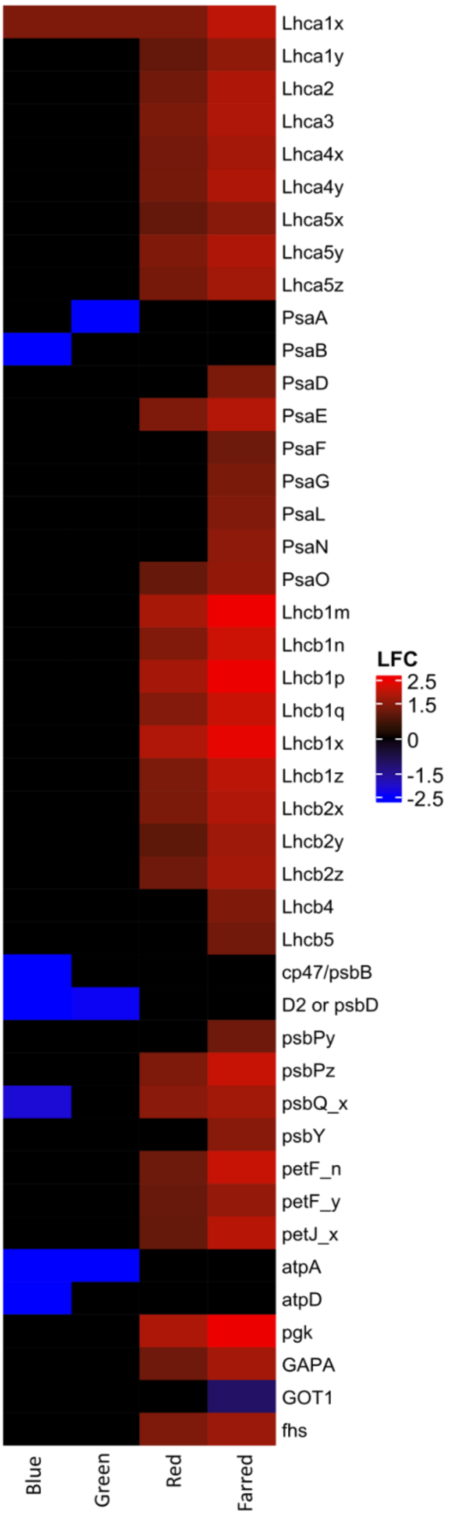

**Figure S17.** Heatmap of photosynthesis, and central carbohydrate metabolism genes of treatment vs White light at day10.

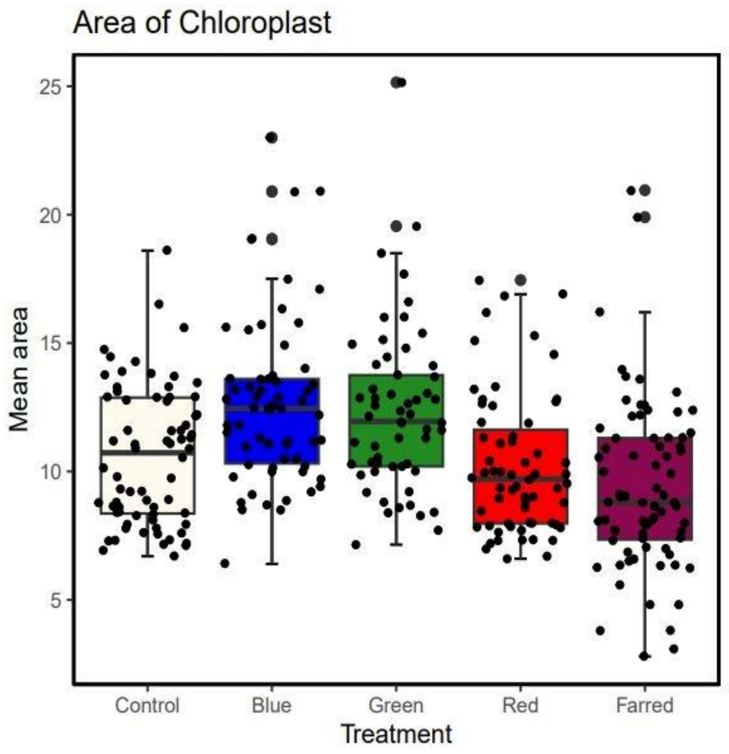

**Figure** S18. Mean area of per chloroplast in box plot.

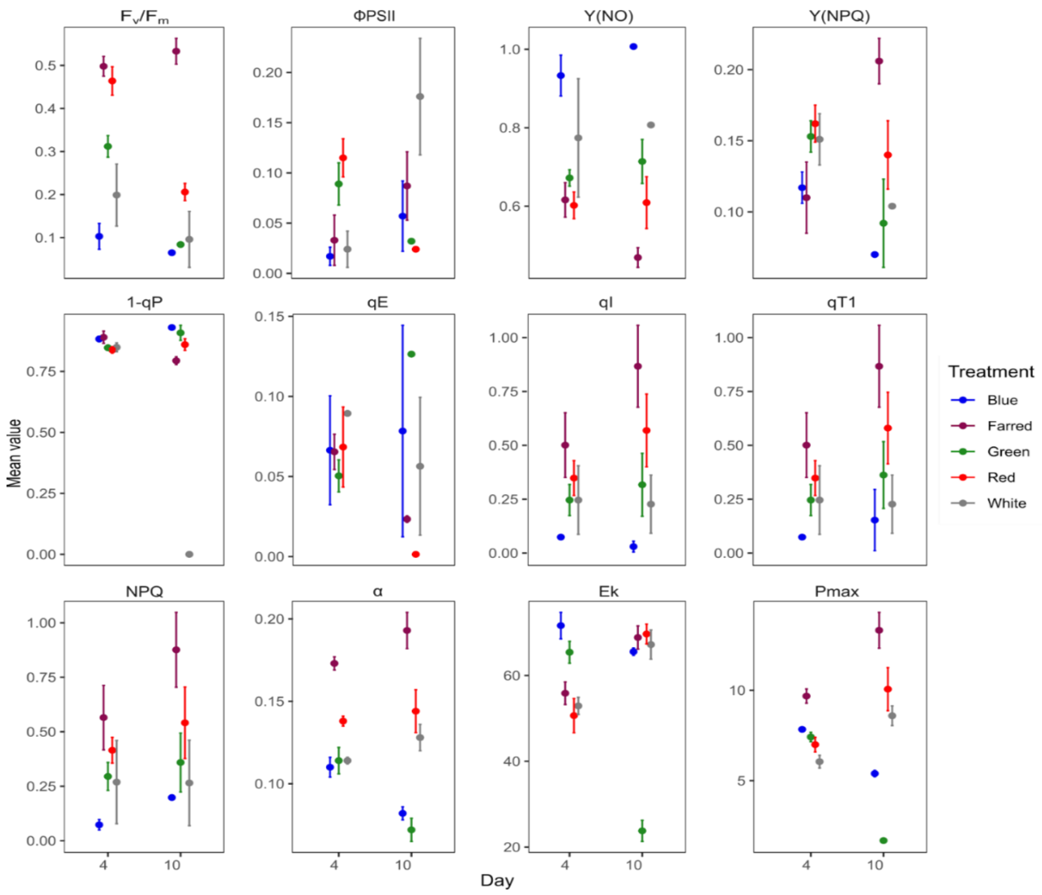

**Figure S19.** Chlorophyll *a* fluorescence based photosynthetic parameter.

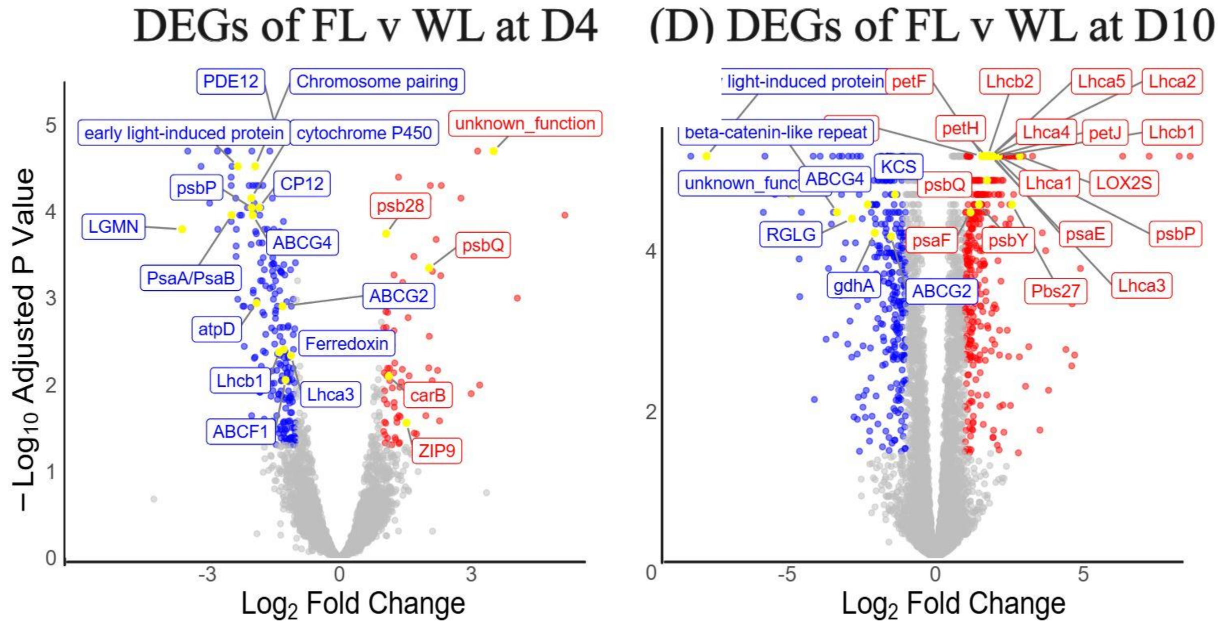

**Figure S20.** Volcano plot showing gene expression of Far-red light (FL) vs. White light (WL), at day4, and at day 10.

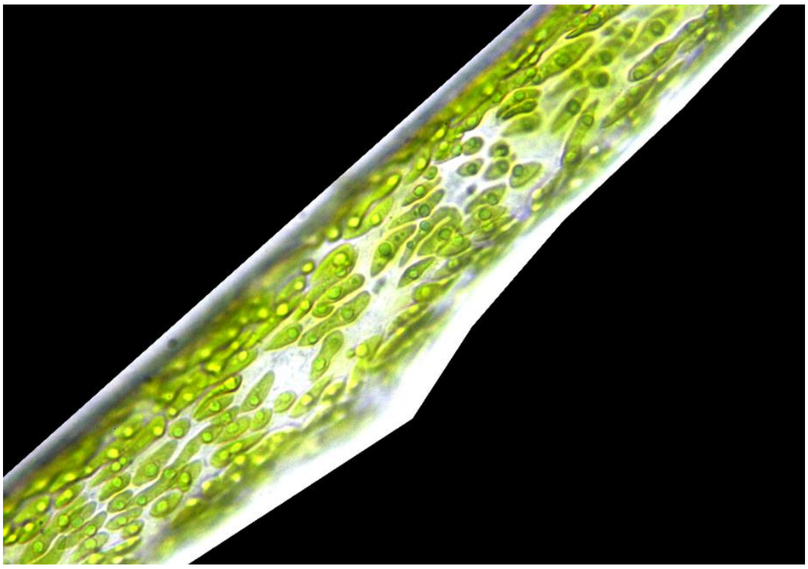

**Figure S21.** Growing algal filament at day10 under far-red light condition.

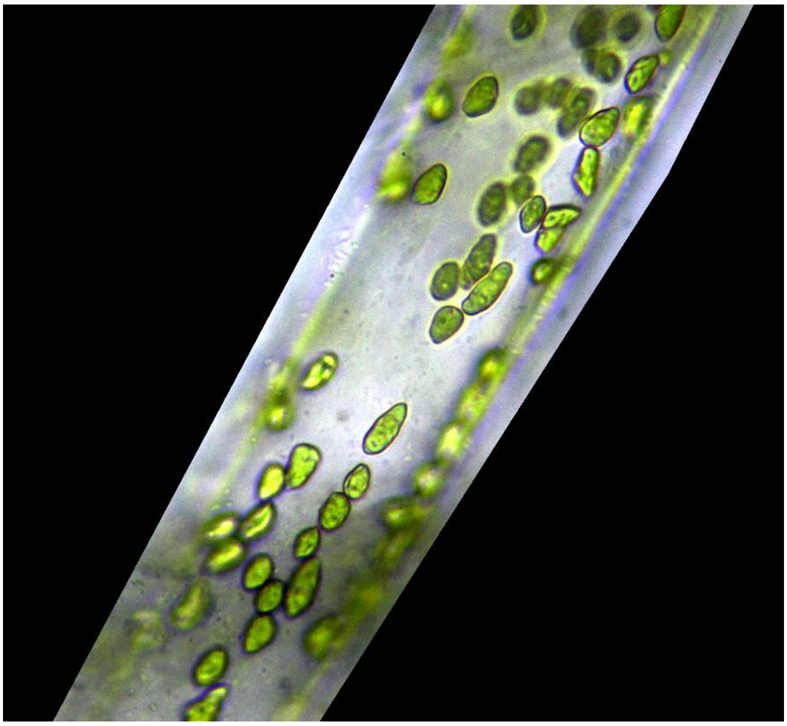

**Figure S22.** Growing algal filament at day10 under red light condition.

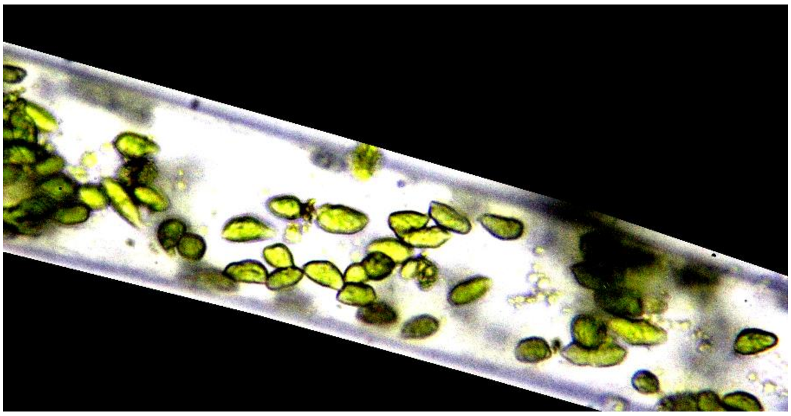

**Figure S23.** Growing algal filament at day10 under white light condition.

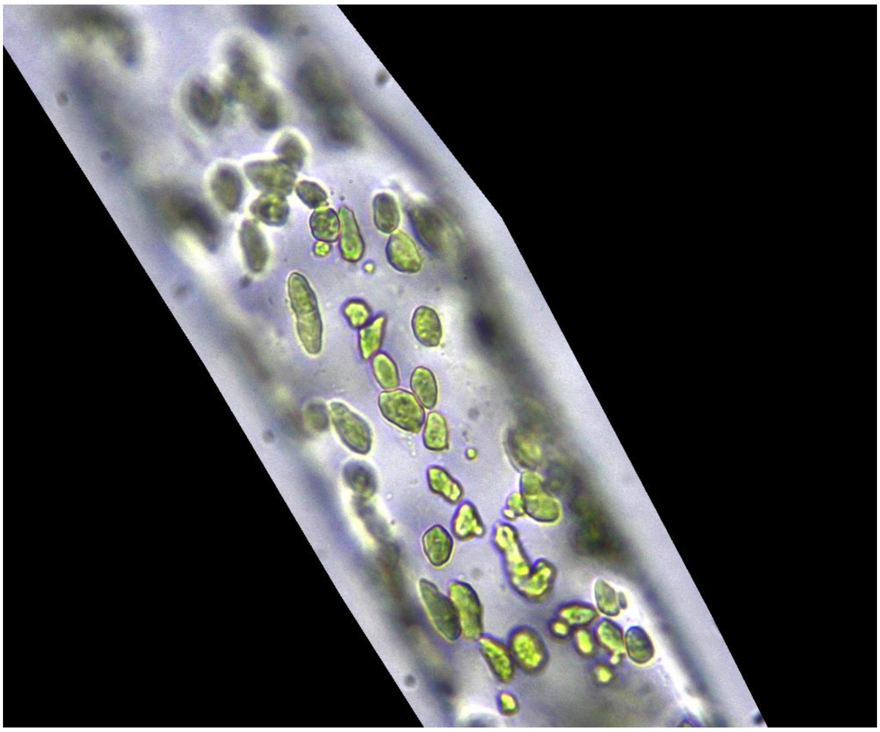

**Figure S24.** Growing algal filament at day10 under blue light condition.

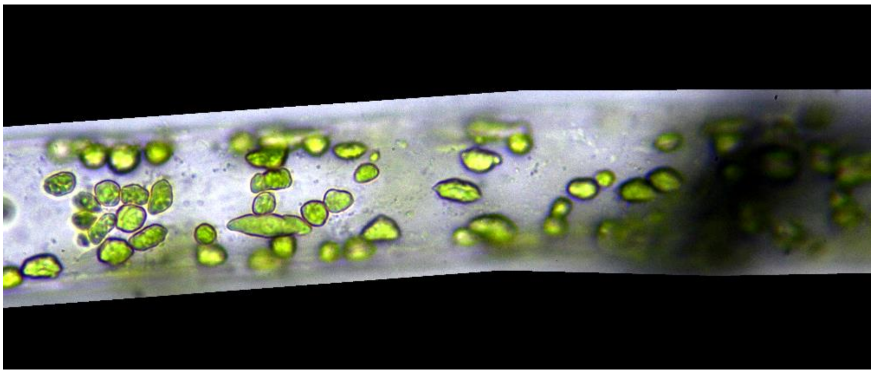

**Figure S25.** Growing algal filament at day10 under green light condition.
